# Reorganization of postmitotic neuronal chromatin accessibility for maturation of serotonergic identity

**DOI:** 10.1101/2021.10.13.463782

**Authors:** Xinrui L. Zhang, W. Clay Spencer, Nobuko Tabuchi, Meagan M. Kitt, Evan S. Deneris

## Abstract

Maturation of transcriptomes encoding unique neuronal identities requires selective accessibility of transcription factors to cis-regulatory sequences in nucleosome-embedded postmitotic chromatin. Yet the mechanisms controlling postmitotic neuronal chromatin accessibility are poorly understood. We used ATAC-seq, ChIPmentation, and single-cell analyses to show that heterogeneous chromatin landscapes are established early and reveal the regulatory programs driving subtype identities of *Pet1*-lineage neurons that generate serotonin (5-HT) neurons. Distal enhancer accessibility is highly dynamic as *Pet1* neurons mature, suggesting the existence of regulatory factors that reorganize postmitotic neuronal chromatin. We find that Pet1 and Lmx1b control chromatin accessibility to select *Pet1*-lineage specific enhancers for 5-HT neurotransmission and synaptogenesis. Additionally, these factors are required to maintain chromatin accessibility during early maturation suggesting that postmitotic neuronal open chromatin is unstable and requires continuous regulatory input. Together our findings reveal postmitotic transcription factors that reorganize accessible chromatin for neuron specialization.

## INTRODUCTION

Specialized neuronal identities are acquired through the differential expression of genes that specify transmitter usage, build dendritic and axonal connectivity, and control synapse formation (Hobert, 2011). The maturation of neuronal functions is an extended process that plays out over several weeks of mid fetal to early postnatal life in mice and through the first few years of life in humans (Stiles and Jernigan, 2010), as postmitotic neuronal transcriptomes mature to a stable state. Unique combinations of sequence-specific transcription factors selectively activate and repress genes to assemble the transcriptomes that encode individual neuron identities (Polleux et al., 2007). Genes with variants associated with autism and other neurodevelopmental disorders exhibit enriched expression in maturing neurons and are often involved in chromatin remodeling and transcriptional regulation; their developmental dysfunction is thought to contribute to disease pathogenesis (De Rubeis et al., 2014; Heavner and Smith, 2020).

Selective expression of neuronal identity genes depends on selective accessibility of promoter and enhancer sequences embedded in postmitotic nucleosomal-organized chromatin to regulatory factor inputs (Gallegos et al., 2018; Perino and Veenstra, 2016). Chromatin accessibility is dynamic as brain regions (Gorkin et al., 2020; de la Torre-Ubieta et al., 2018) and neurons (Frank et al., 2015; Preissl et al., 2018; Stroud et al., 2020; Trevino et al., 2021) mature. Yet, we have a poor understanding of how the dynamics of accessible chromatin in postmitotic neurons prefigures acquisition of neurotransmitter identities and myriad other molecular and morphological features of specialized neurons. Further, the regulatory factors involved in selecting postmitotic accessible chromatin regions to enable sequence-specific activation of unique combinations of identity features have not been identified.

Control of serotonin (5-HT) neuron development is of particular interest as 5-HT has expansive modulatory effects on central neural circuitry (Celada et al., 2013). Altered serotonergic gene expression, brought about by either genetic or environmental factors, has been implicated in neuropsychiatric disorders including depression, stress related anxiety disorders, autism, obsessive compulsive disorder, and schizophrenia (Deneris and Wyler, 2012). Postmitotic 5-HT neuron precursors terminally differentiate in the ventral hindbrain to acquire the capacity for 5-HT synthesis, reuptake, vesicular transport, and degradation through the coordinate expression of the 5-HT neurotransmission genes *Tph2*, *Ddc*, *Gch1*, *Slc6a4* and *Slc22a3*, *Slc18a2*, and *Maoa/b* (Deneris and Gaspar, 2018). Subsequently, newborn 5-HT neurons enter a maturation stage during which serotonergic transcriptomes are highly dynamic as 5-HT neurons refine their terminal identities, migrate to populate the various raphe nuclei, acquire mature firing properties, and establish expansive connectivity throughout the brain and spinal cord (Deneris and Gaspar, 2018). Little is known about the regulatory mechanisms that underlie the postmitotic maturation of neurons.

The ETS domain factor Pet1 and the LIM HD factor Lmx1b are terminal selector transcription factors (TFs) in 5-HT neurons as they are continuously expressed beginning at the postmitotic precursor stage, control their own expression through direct positive autoregulation, and are required for the acquisition of 5-HT neuron terminal identity through sequence-specific transcriptional activation of 5-HT pathway genes (Deneris and Gaspar, 2018; Hobert, 2008). Pet1 and Lmx1b are also required for the acquisition of serotonergic firing characteristics and formation of long-distance profusely arborized 5-HT axon architectures throughout the brain and spinal cord (Donovan et al., 2019; Liu et al., 2010; Wyler et al., 2016). Pet1’s function in 5-HT neurons is clinically significant as recent studies identified biallelic loss of function of *FEV*, the human ortholog of *Pet1*, in autism cases suggesting that defects in transcriptional control of 5-HT neurons are a potential path to human neurodevelopmental disorders (Doan et al., 2019).

The low abundance of many postmitotic neuron-types has hindered the assay of their accessible chromatin landscapes and cis-regulatory architectures. Here, we adapted bulk and single cell ATAC-seq, scRNA-seq, and ChIPmentation protocols to map accessible chromatin and cis-regulatory elements of developing 5-HT neurons. We then used these combined maps along with Pet1 and Lmx1b knockout mouse models and occupancy mapping to identify a critical role for these TFs in directly regulating the chromatin accessibility of 5-HT neurotransmission and synapse genes. Our findings show that Pet1 and Lmx1b control neuron maturation not only through sequence-specific activation of select terminal effector genes but also by reorganizing postmitotic accessible chromatin at cis-regulatory elements (CREs). Lastly, analysis of Pet1 and Lmx1b regulated CREs reveal a novel function of these TFs in 5-HT synaptogenesis.

## RESULTS

### The unique accessible chromatin landscape of maturing postmitotic *Pet1* neurons

5-HT neurons are generated from *Pet1*+ postmitotic precursors located in the embryonic ventral hindbrain from the isthmus to the caudal region of the myelencephalon (Hendricks et al., 1999). Therefore we used the *Pet1::EYFP* transgenic mouse line to flow sort E14.5 Yfp+ neurons (*Pet1* neurons) from the entire longitudinal extent of the hindbrain for assay of accessible chromatin in maturing 5-HT neurons (Scott et al., 2005a, 2005b) (Supplemental Fig 1a-c). We generated three ATAC-seq biological replicates with a mean of 29M unique paired-end mapped reads per sample (Supplemental Fig 1d-g) and identified 68,871 peaks of transposase accessible chromatin (TAC) at E14.5, and 59,323 TACs in an equal number of control Yfp- (*Pet1*^neg^) cells collected from the same hindbrain regions at the same developmental age, which comprise diverse non-serotonergic cell types. Principal component analysis (PCA) showed that cell type identity accounted for 93% of the variance when comparing *Pet1* neuron ATAC-seq replicates with *Pet1*^neg^ ATAC-seq samples and with previously published ATAC-seq datasets obtained from bulk tissue dissections (Fig 1a). The accessible chromatin landscape of *Pet1* neurons is distinct from previously published bulk hindbrain ATAC-seq (Gorkin et al., 2020), which indicates that bulk analyses did not capture the chromatin accessibility profiles of low abundance *Pet1* neurons (Fig 1a). Indeed, of the TACs we identified in *Pet1* neurons at E14.5, 31% are unique and not found in *Pet1*^neg^ cells of mouse hindbrain and 22% are not previously identified in any other cell type (Gorkin et al., 2020) (Fig 1b). The large number of unique TACs may therefore more precisely define *Pet1* neuron identity than gene expression.

**Figure 1:**
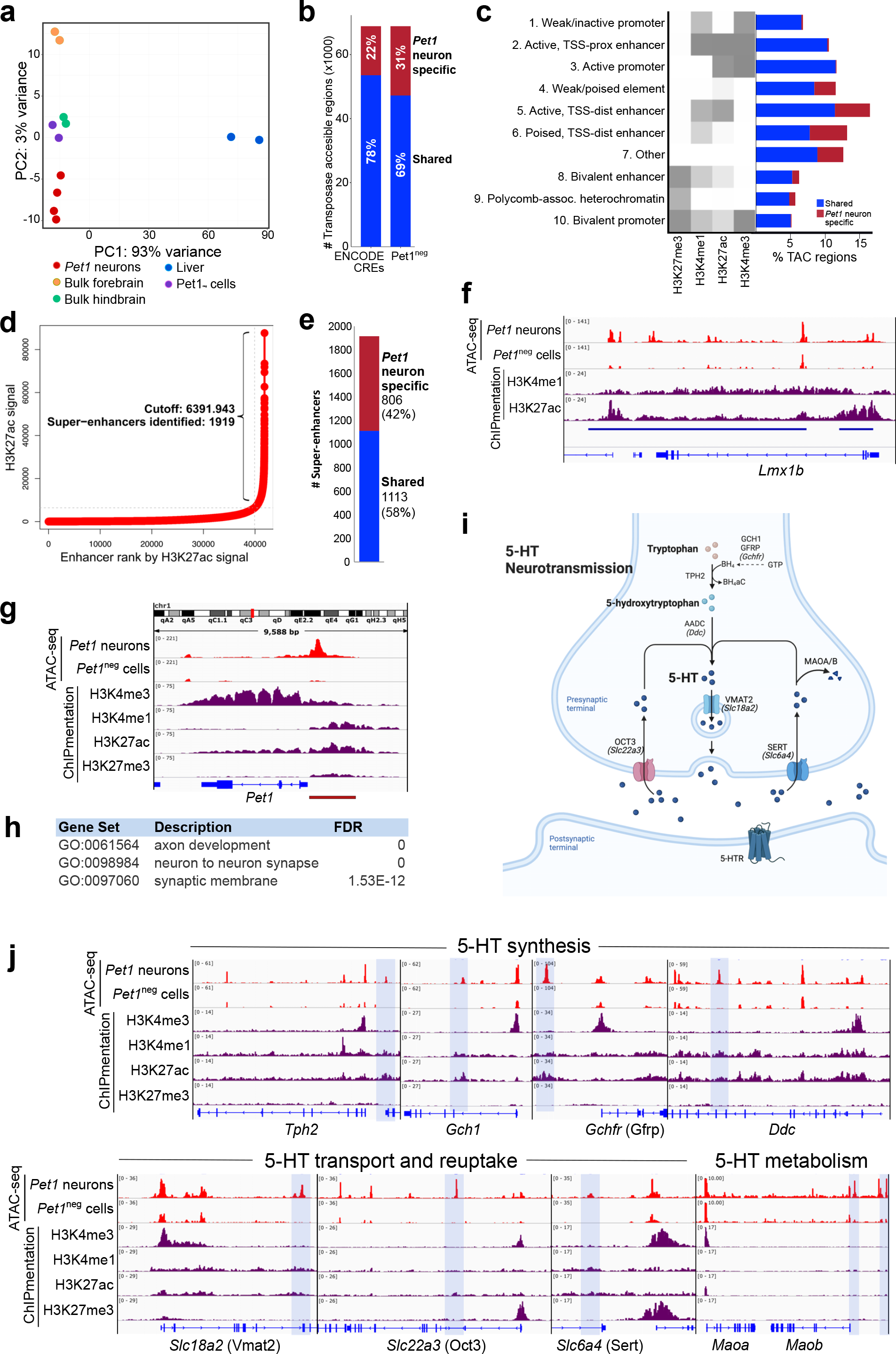
Unique distal enhancers and super-enhancers define the *Pet1* lineage. (a) PCA reveals the unique accessible chromatin landscape of *Pet1* neurons. Dots of the same color represent biological replicates. (b) Bar plot showing percentage of unique ATAC-seq peaks (transposase accessible chromatin “TAC” regions) from E14.5 *Pet1* neurons when compared to CREs annotated by ENCODE datasets (Gorkin et al., 2020) (left bar) or with TACs in E14.5 *Pet1*^neg^ cells from the same hindbrain region (right bar). (c) Emission probabilities for histone modifications in ten ChromHMM states and the percent genomic coverage of each chromatin state for E14.5 *Pet1* neuron specific TACs and *Pet1* neuron TACs shared with other tissues in ENCODE ATAC-seq datasets. (d) Distribution of H3K27ac signal across all enhancers identified in E14.5 *Pet1* neurons. Enhancers with exceptionally high read counts (bracket) were designated super-enhancers. (e) Bar plot of *Pet1* neuron super-enhancers that are unique to *Pet1* neurons or overlap (“shared”) with annotated enhancers from ENCODE datasets. (f) Genome browser view of the 5-HT neuron specific super-enhancer across the *Lmx1b* locus defined as an extended stretch of H3K27ac and H3K4me1 enrichment. (g) Genome browser views of ATAC-seq and ChIPmentation signals at the *Pet1/Fev* locus. Red bar denotes the conserved *Pet1* enhancer region that is sufficient to drive 5- HT neuron-specific transgene expression (Scott et al., 2005a, 2005b). (h) Go ontology of Pet1 neuron specific TACs (not found in other tissues in ENCODE ATAC-seq datasets) (i) Diagram of 5-HT neurotransmission. (j) Genome browser views of ATAC-seq and ChIPmentation peaks at 5-HT neurotransmission genes. Light blue shading highlights *Pet1* neuron specific TACs.

To determine the functions of 5-HT neuron TACs, we next used ChIPmentation (Schmidl et al., 2015) to profile the genome-wide distribution of histone posttranslational modifications associated with transcriptional activation or repression in flow sorted E14.5 *Pet1* neurons. With the segmentation algorithm ChromHMM (Ernst and Kellis, 2017), we defined ten chromatin states by assessing the combinatorial pattern of histone marks at each genomic locus (Fig 1c; Supplemental Fig 1h). Active and poised distal enhancer regions are over-represented among the *Pet1* neuron specific TACs (Fig 1c; Supplemental Fig 1i-j). Our ChIPmentation analysis also identified 1919 genomic regions that fulfill the criteria for super-enhancers based on the presence of unusually expansive H3K27ac signals (Fig 1d) (Lovén et al., 2013; Whyte et al., 2013). Comparing these to previously annotated enhancers in other cell types (Gorkin et al., 2020), we found that a significant percentage (42%) of the super-enhancers we detected in *Pet1* neurons were not detected in bulk mouse forebrain, hindbrain, or liver tissues (Fig 1e), suggesting that they are highly specific to the *Pet1* lineage. As an example, a 82.9kb super-enhancer detected in *Pet1* neurons but not *Pet1*^neg^ neurons spans the *Lmx1b* locus (Fig 1f).

Notably, a distinct TAC was found immediately upstream of the *Pet1* promoter in *Pet1* neurons but not *Pet1*^neg^ cells (Fig 1g). This TAC shows enrichment of H3K4me1 and H3K27ac marks suggesting the presence of an active enhancer (Fig 1g). Indeed, previous transgenic assays, *in vivo*, demonstrated that this upstream region directs robust reporter expression to developing and adult 5-HT neurons present in each of the midbrain, pontine, and medullary raphe nuclei (Krueger and Deneris, 2008; Scott et al., 2005a, 2005b), hence providing functional validation of serotonergic enhancers we have defined with ATAC-seq and ChIPmentation. Importantly, genes associated with *Pet1* neuron specific TACs are enriched for axon and synapse functions (Fig 1h). Indeed, we found Pet1 neuron specific TACs for 5-HT neurotransmission genes (Fig 1i-j). Together, our ATAC-seq and ChIPmentation analyses define the open chromatin landscape of maturing 5-HT neurons and reveal that predominantly distal CREs implicated in axon and synapse related functions define the *Pet1* postmitotic lineage.

### Heterogeneous distal enhancer accessibility defines *Pet1* neuron subtypes

Adult 5-HT neurons possess diverse transcriptomes that encode functional subtypes (Okaty et al., 2015, 2020; Ren et al., 2019; Wylie et al., 2010), yet the developmental mechanisms underlying 5-HT neuron heterogeneity are not understood. To determine whether 5-HT neuron heterogeneity is coded early in accessible chromatin, we carried out single cell ATAC-seq (scATAC-seq) with E14.5 flow sorted *Pet1* neurons, which were processed using the 10x Genomics Chromium microfluidic droplet scATAC-seq method. The resulting library was sequenced to a depth sufficient to yield >50,000 paired-end reads per nucleus. After filtering based on standard quality control measures (Supplemental Fig 2a), we retained 1,692 nuclei for analysis. This led to the identification of 124,111 TACs, 60,117 of which were not identified in our bulk *Pet1* neuron ATAC-seq (Fig 2a). We observed high concordance (Spearman’s correlation = 0.9) between our pseudo-bulk scATAC-seq dataset and our bulk ATAC-seq data as illustrated by the highly similar transposase accessible chromatin patterns in the *Pet1* upstream enhancer region (Fig 2b).

**Figure 2:**
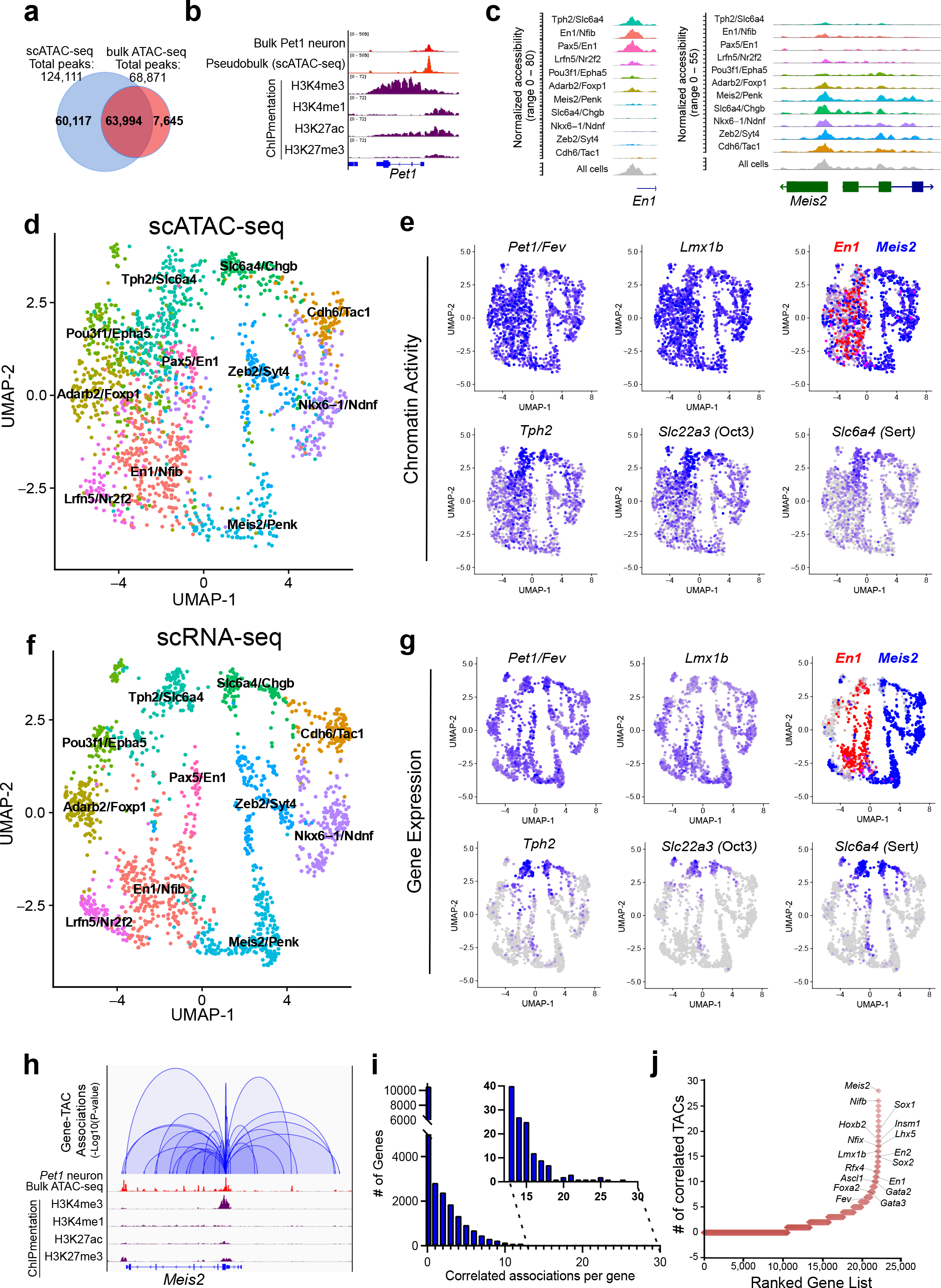
Heterogeneity in chromatin accessibility and transcriptomes reveals transcriptional programs driving *Pet1* neuron subtype identities. (a) Venn diagram showing overlap of single cell *Pet1* neuron TACs versus bulk ATAC-seq TACs. (b) Genome browser view of *Pet1* locus showing high concordance between the pseudobulk scATAC-seq and bulk FACS-purified *Pet1* neuron ATAC-seq at E14.5. (c) scATAC-seq tracks showing the aggregated chromatin accessibility peaks of *En1* (left) and *Meis2* (right) for each cluster. (d) UMAP visualization of chromatin accessibility of single E14.5 *Pet1*-lineage neurons. scATAC-seq is projected onto scRNA-seq UMAP space. Cells are colored by scATAC-seq cluster assignment. Genes associated with the top two marker TACs for each cluster are labeled. (e) The chromatin accessibility profiles of select genes across UMAP visualized scATAC-seq clusters. (f) UMAP visualization of single E14.5 *Pet1* neurons based on gene expression. Cells are colored by scRNA-seq cluster assignment. (g) The expression profile of example 5-HT neuron identity genes across all scRNA-seq clusters in UMAP. (h) Representative gene with high number of associated TACs. Loops represent statistically significant TAC-gene association. Loop height represents the p-value of TAC-gene correlation. (i) Number of TAC-gene associations for all genes. (j) Number of significantly correlated TACs for each gene. Genes are ranked by the numbers of associated peaks. Select TFs with known or potential functions in the *Pet1* lineage are labeled.

We grouped single *Pet1* neurons to identify 11 major subtypes with unique chromatin accessibility patterns (Fig 2d; Supplemental Fig 2b). TACs within *Pet1* and *Lmx1b* loci are robustly accessible in virtually all cells within the 11 identified subtypes (Fig 2e). Like *Pet1* and *Lmx1b*, two-thirds (63%) of TACs are broadly accessible among multiple subtypes, including TACs of 5-HT neurotransmission genes *Tph2*, *Slc22a3*, and *Slc6a4* (Fig 2e; Supplemental Fig 2c). In contrast, one-third of TACs (36.6%) are unique to a single cluster. Subtype-specific TACs are predominantly distal gene enhancers as previously defined by chromatin state (Fig 1c; Supplemental Fig 2d-e) and are often observed near genes that mark known 5-HT neuron subdivisions (Supplemental Fig 2b). For example, transcription factor En1 is enriched in the rostral rhombencephalic 5-HT neurons, whereas Hox gene cofactor Meis2 is selectively expressed in the caudal rhombencephalic 5-HT neurons (Wylie et al., 2010). Six of the scATAC-seq clusters share an unique TAC region immediately upstream of the TSS of the rostral marker gene *En1* while the remaining five clusters have more accessible TAC regions near the promoter of the caudal 5-HT neuron marker gene *Meis2* (Fig 2c-e). Thus, our data demonstrate that diverse single cell chromatin landscapes are established early in postmitotic *Pet1* neurons and likely prefigure transcriptomic and functional heterogeneity of adult 5-HT neurons.

To analyze the target genes of subtype-specific TACs, we performed scRNA-seq in parallel using 10x Genomics and E14.5 flow sorted *Pet1* neurons (Supplemental Fig 2f-g). We obtained over 80,000 reads per cell for 1,612 cells, representing a median of 4,010 genes per cell and a total of 19,707 genes across all cells. 95% of the flow sorted *Pet1* neurons robustly expressed both *Pet1* and Yfp, validating that our cell sorting method yields relatively pure populations of *Pet1* neurons. The small numbers of Yfp- cells were removed from further analysis. We next co-embedded scATAC-seq with scRNA-seq data, then linked TACs to a gene if logistic regression detected a statistical association between gene expression and the binarized chromatin accessibility at a genomic region within 500kb (Fang et al., 2021). Applying this method to our single cell datasets yielded a total of 41,191 enhancer-promoter pairings (Supplemental Table). Each gene is on average linked to only 1.8 TACs and most genes are associated with either none or a single TAC (Fig 2i). However, 10% of genes are associated with >5 TAC regions, and a select group of genes (0.045%) are associated with >20 TACs (Fig 2h-j; Supplemental Fig 3a). Gene ontology (GO) analysis of the top 1,500 genes with the highest number of linked CREs revealed that they are highly enriched for genes involved in brain development (Supplemental Fig 3b), suggesting that a large number of CREs converge to fine-tune or safeguard the transcription of key neuronal regulators (Ma et al., 2020; Trevino et al., 2021). Among these genes are known TFs of the 5-HT neuron gene regulatory network such as Pet1 and Lmx1b (Fig 2j).

We identified many subtype-specific TFs constituting a potential transcription factor code that underlies 5-HT neuron heterogeneity (Supplemental Fig 3c-d). For example, Nkx6-1 activity, gene expression, and chromatin accessibility are enriched in the Nkx6-1/Ndnf cluster (Supplemental Fig 3c-d). TFs expressed in a *Pet1* neuron subtype have gene regions that are also more accessible in that subtype (Supplemental Fig 3d). Overall most subtype-specific TACs are correlated with the subtype-specific expression pattern of their linked genes; however, we noticed that the expression of 5- HT identity genes *Tph2*, *Slc6a4*, and *Slc22a3* are low or undetectable in most *Pet1* neurons at E14.5 despite their relatively uniform accessible chromatin in *Pet1* neurons at this stage (Figs 2e and 2g). The vast majority of *Pet1*-lineage neurons robustly express these 5-HT identity genes in the adult mouse brain (Supplemental Fig 3e) (Donovan et al., 2019; Okaty et al., 2015, 2019; Ren et al., 2019). Immunofluorescence analysis of Tph2 protein levels showed that Tph2+ neurons emerge in the highest fractions in *Pet1* neurons of the dorsal raphe where 5-HT neurons are first born and have progressed further in their maturation than those in other raphe nuclei (Supplemental Fig 3f). This suggests that chromatin accessibility may be remodeled early during maturation to prime neuron identity before the transcription of terminal effector genes.

### A highly dynamic stage of postmitotic accessible chromatin reorganization

Previous whole genome RNA-seq study revealed that after their postmitotic birth and acquisition of 5-HT-type transmitter identity, 5-HT neurons undergo a prolonged maturation process that involves the expression of genes needed for the fully mature, functional 5-HT phenotypic state (Wyler et al., 2016). One possibility is that Pet1 neurons are born with fully mature chromatin that is accessible to all regulatory factors needed for the eventual transcription of 5-HT genes. In this case, developmental gene trajectory is likely supported by the maturation of regulatory factor interactions. Another possibility is that *Pet1* neurons are born with immature chromatin and the opening of new TACs at various loci would be a fundamental step in controlling *Pet1* neuron gene expression and developmental specialization.

To investigate whether chromatin accessibility in maturing *Pet1* neurons is developmentally dynamic, we compared TACs at E11.5 when postmitotic 5-HT neuron precursors are abundant, E14.5 when *Pet1* neurons are actively extending dendrites and axons, and E17.5 when *Pet1* neurons are coalescing to form the raphe nuclei and innervating distal target brain regions (Donovan et al., 2019) (Supplemental Fig 4a-b).

PCA showed high concordance of ATAC-seq replicates within each time point with developmental state accounting for the majority of variance (Fig 3a). Significant gains and losses in chromatin accessibility of TSS-distal TACs occurred throughout embryonic maturation (Fig 3b-d; Supplemental Fig 4c-d,i). Of the TACs detected in *Pet1* neurons, 16,493 “open” (gain accessibility; fold change>2 and FDR <0.01) between E11.5 and E14.5, while 4,148 open from E14.5 to E17.5 (Fig 3c). Concurrently, 9,402 E11.5 TACs “close” (lose accessibility, fold change>2 and FDR<0.01) by E14.5, and 1504 TACs close between E14.5 and E17.5 (Fig 3c). This suggests that chromatin remodeling is highly dynamic immediately following cell cycle exit and then gradually stabilizes. Assessment of chromatin states showed that active enhancers are over-represented among TACs opening between E11.5 and E17.5 (Fig 3b; Supplemental Fig 4e-g). *Pet1* neuron super-enhancer accessibility is particularly highly dynamic during postmitotic neuron maturation, with the vast majority (93%) of TACs either gaining or losing accessibility between E11.5 and E17.5 (Supplemental Fig 4h).

**Figure 3:**
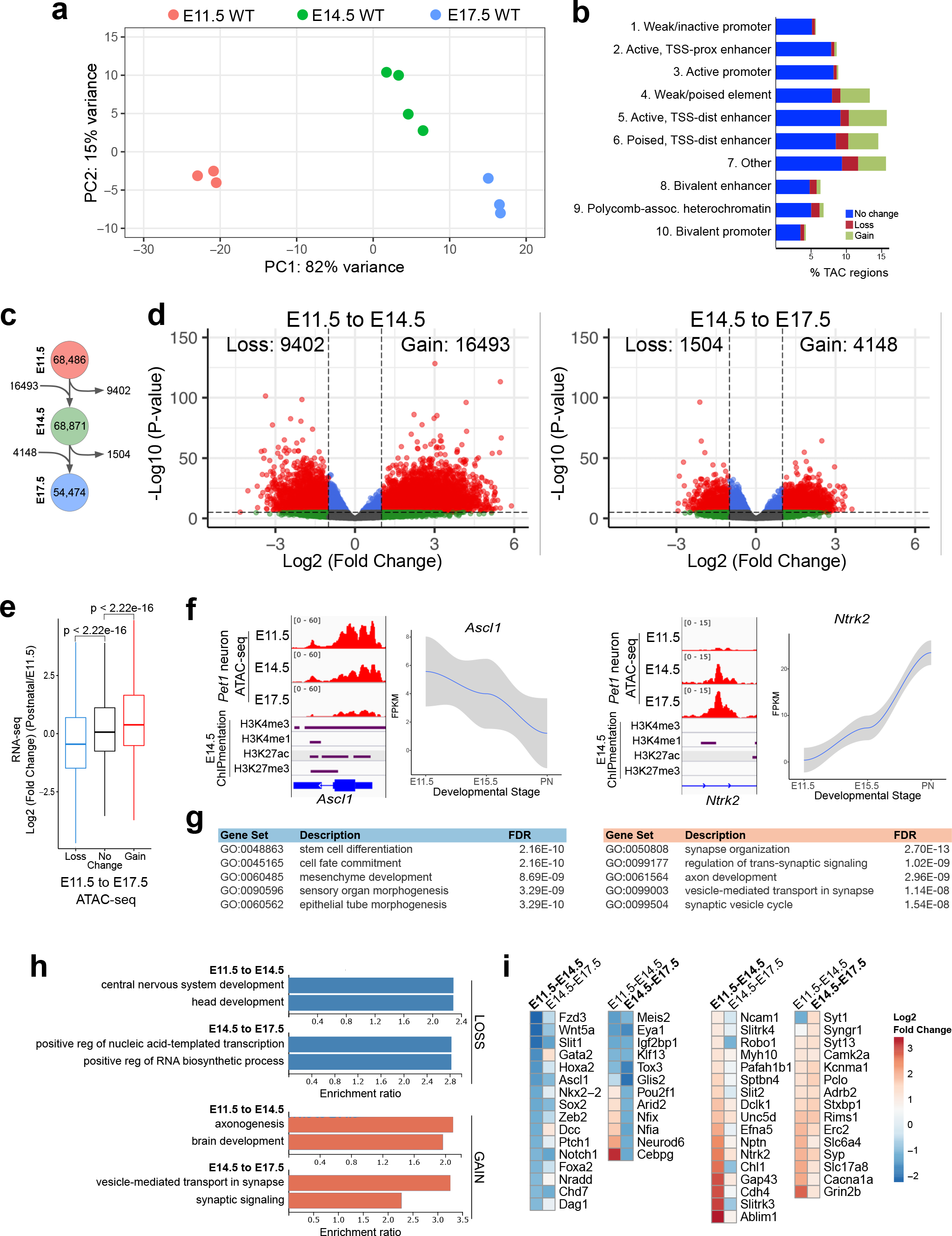
Gene regulatory dynamics driving *Pet1* neuron maturation. (a) PCA showing the distribution of *Pet1* neuron ATAC-seq data over two principal components. (b) Distribution of chromatin states that exhibit gain (green), loss (red), or no change (blue) in chromatin accessibility from E11.5 to E17.5. (c) Number of TACs detected at each developmental time point (circles) and the number of TACs that are gained, lost, or propagated to the next time point (arrows). (d) Volcano plots showing the differential chromatin accessibility of TACs between consecutive time points, with fold change cutoff of 2-fold and significance cutoff of FDR <0.01. Each dot represents one ATAC-seq peak (TAC). (e) Box plots showing the distribution of the log2 expression values of the linked genes for TACs that exhibit loss (blue), no change (black), or gain (red) in accessibility between E11.5 and E17.5. Significance of accessibility to gene expression correlation is calculated by Wilcoxon ranks sum test. (f) Genome browser view of ATAC-seq and E14.5 ChIPmentation signals at representative genes that decrease (e.g. *Ascl1*) or increase (e.g. *Ntrk2*) in accessibility between E11.5 and E17.5. Parallel trends are seen in gene expression trajectories (panels on the right of tracks). (g) Top five gene ontology biological process enrichment terms for genes linked to TACs that lose (left) or gain (right) accessibility from E11.5 to E17.5. (h) Bar graphs showing the enrichment ratio of the top gene ontology biological process enrichment terms for genes linked to TACs that lose (blue) or gain (orange) accessibility from E11.5 to E14.5 and from E14.5 to E17.5. (i) Heatmap showing the log 2 fold-change in chromatin accessibility for TACs of select genes associated with the top GO biological processes of each developmental stage

Next we compared our bulk ATAC-seq datasets at E11.5, E14.5, and E17.5 with our previously published bulk 5-HT neuron RNA-seq data obtained at E11.5, E15.5, and postnatal day 2-3 (Wyler et al., 2016). Using our TAC-to-gene assignments (Fig 2i-j), we found that changes in chromatin accessibility at cis-regulatory regions are significantly correlated with the expression trajectories of their target genes (Fig 3e-f). GO analysis revealed that genes associated with closing TACs are important for neural precursor functions, whereas the genes linked to opening TACs are strongly enriched for axon development between E11.5 and E14.5 and synaptic signaling between E14.5 and E17.5 (Fig 3g-i). Dynamic TACs are highly enriched for diverse TF binding motifs, suggesting that combinatorial TF activities contribute to *Pet1* neuron maturation (Supplemental Fig 4i). Together these findings support a model in which the dynamic remodeling of serotonergic CRE accessibility drives gene expression trajectories required for maturation of 5-HT identity while shutting down expression of genes required at earlier stages of development.

### Pet1 reorganizes chromatin accessibility of 5-HT neurotransmission genes

The highly dynamic reorganization of TACs in maturing postmitotic *Pet1* neurons predicts the existence of regulatory factors that shape postmitotic neuron chromatin landscapes. However, such factors have not been described. Notably, TACs that are unique to *Pet1*-lineage neurons are enriched for ETS-factor binding sites (Supplemental Fig 5a). Thus, we hypothesized that Pet1 controls acquisition of 5-HT identity by controlling chromatin accessibility. *Pet1-/-* 5-HT neurons are an ideal model with which to examine transcription factor function in shaping the chromatin, as these neurons are retained in the brain in normal numbers and *Pet1::EYFP* transgene expression in these cells is robustly retained at fetal stages to permit their isolation and analysis (Wyler et al., 2016). PCA of ATAC-seq datasets obtained from *Pet1* neurons from *Pet1::EYFP+, Pet1-/-* hindbrain tissue at E11.5, E14.5, and E17.5 (Supplemental Fig 5b-c) showed that *Pet1-/- Pet1-*lineage neurons (*Pet1-/-* neurons) cluster closely with wildtype *Pet1* neurons (Fig 4a), consistent with our observation that Pet1-deficient 5-HT neurons do not adopt another neuron-type identity. However, we found a striking overall reduction of Pet1 footprints and decreased accessibility of many TSS-distal TACs in *Pet1-/-* neurons compared to wildtype neurons at all three embryonic stages (Fig 4b-c; Supplemental Fig 5d). To determine whether Pet1 directly controls chromatin accessibility in these chromatin regions, we performed CUT&RUN with E14.5 flow sorted *Pet1* neurons to generate a genomic occupancy map for Pet1. The CUT&RUN map significantly overlapped with our previous ChIP-seq of Pet1 but identified occupancy at several additional 5-HT neurotransmission genes whose expression is reduced in Pet1-deficient 5-HT neurons (Wyler et al., 2016) (Supplemental Fig 5e). CUT&RUN data showed that Pet1 directly occupies at least 53% of Pet1-dependent TACs (Fig 4d-e). Moreover, 73% of all Pet1-regulated TACs and 80% of Pet1-regulated TACs with Pet1 occupancy are TACs unique to *Pet1* neurons (Fig 4f).

**Figure 4:**
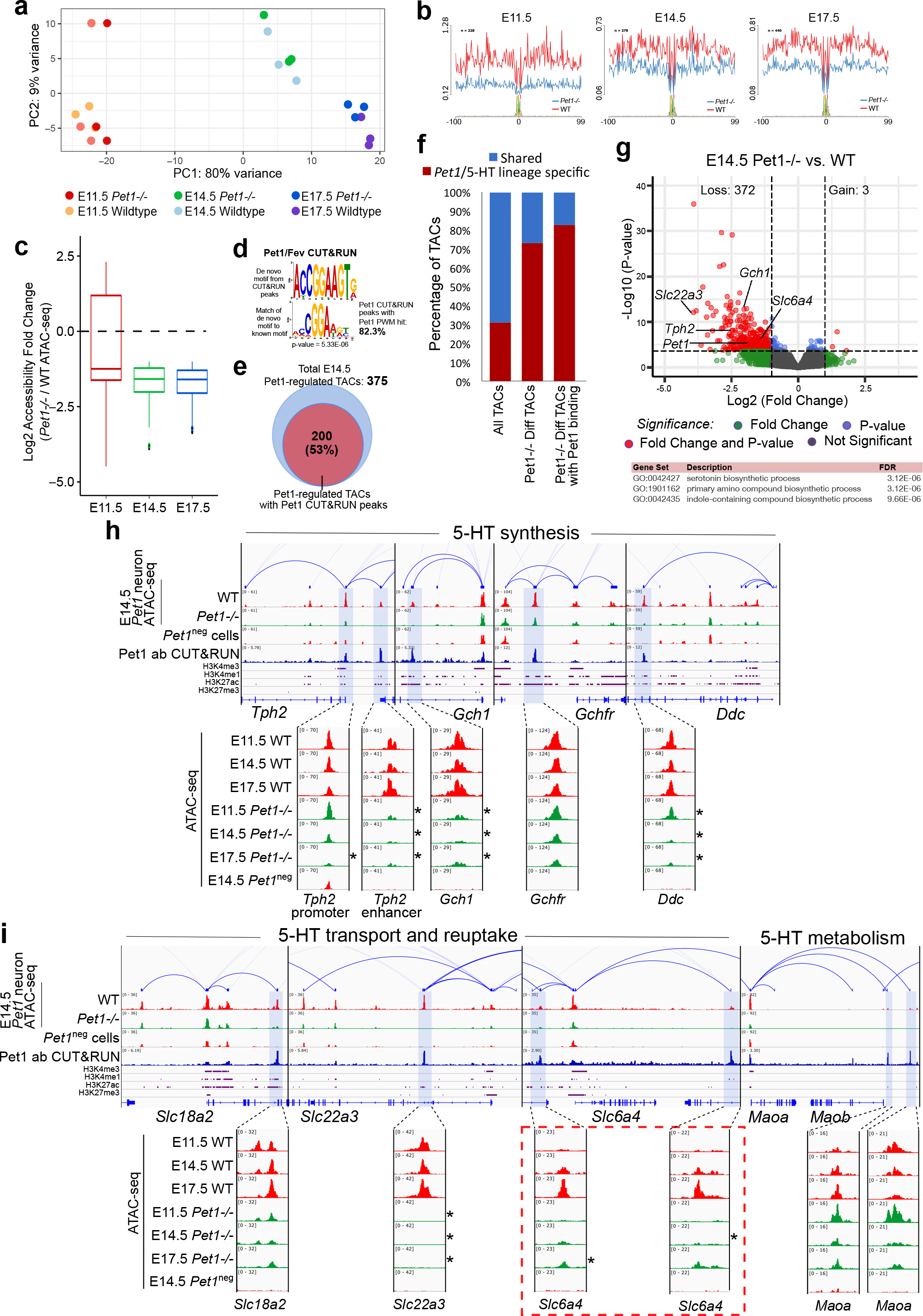
Pet1 reorganizes postmitotic chromatin accessibility. (a) PCA showing the distribution of ATAC-seq data for wildtype and *Pet1-/- Pet1*-lineage neurons (*Pet1-/-* neurons) over two principal components. (b) *In vivo* Pet1 footprints derived from Tn5 insertion frequency (ATAC-seq reads) over representative ETS-factor motifs within *Pet1* neuron accessible chromatin regions. *Pet1-/-* neurons show reduced Pet1 footprints at E11.5 (left), E14.5 (middle), and E17.5 (right) compared to wildtype neurons. (c) Box plots showing the distribution of the log2 fold change in accessibility of all TACs in *Pet1-/-* vs. wildtype *Pet1* neurons at three developmental time points. (d) The top significantly enriched motif within Pet1 CUT&RUN peaks as determined by *de novo* MEME motif analysis significantly matches (p=5.33E-06) the Pet1/FEV high- affinity binding site previously defined *in vitro* (Wei et al., 2010) based on analysis by the TOMTOM Motif Comparison Tool (top panel). (e) The fraction of Pet1-regulated TACs containing Pet1 CUT&RUN peaks. (f) The fraction of all *Pet1* neuron TACs, *Pet1* neuron TACs that are dependent on Pet1, and Pet1-dependent TACs occupied by Pet1 that are specific to *Pet1*/5-HT lineage or shared with *Pet1*^neg^ cells. A significant fraction of Pet1-regulated TACs is unique to *Pet1* neurons and absent from *Pet1*^neg^ cells. (g) Volcano plot showing the differential chromatin accessibility of TACs between *Pet1-/-* and wildtype *Pet1* neurons at E14.5, with fold change cutoff of 2-fold and significance cutoff of FDR <0.01 (top). Highest FDR GO terms for Pet1-dependent TACs (in red) that are occupied by Pet1 show strong enrichment for serotonin biosynthesis. Select genes required for 5-HT synthesis are labeled on the volcano plot. (h-i) Genome browser tracks showing the gene-enhancer pairings and the ATAC-seq, Pet1 CUT&RUN, and histone modification ChIPmentation signals at 5-HT identity genes. Zoomed-in inserts display the changes in chromatin accessibility between wildtype (red) and *Pet1-/-* (green) *Pet1* neurons within the highlighted (blue) regions at three developmental time points. Two TACs of the gene *Slc6a4* (red dashed line box) gain accessibility in wildtype 5-HT neurons from E11.5 to E17.5 but fail to open to the same extent in *Pet1-/-* neurons. Asterisks denote FDR<0.01 and fold change >2 relative to wildtype at the same time point.

GO analysis revealed that the Pet1 occupied TACs that close in *Pet1-/-* neurons are associated with genes required for serotonin synthesis (Fig 4g; Supplemental Fig 5f). Indeed, we found a significant loss of accessibility of enhancers either upstream of the TSS or within introns of *Tph2*, *Gch1*, and *Ddc* (Fig 4h). TACs associated with the serotonin reuptake transporter genes *Slc22a3* and *Slc6a4* are also significantly reduced in accessibility (Fig 4i). Importantly, enhancers that require Pet1 for accessibility are unique to *Pet1* neurons, while the non-*Pet1* neuron specific TACs of 5-HT pathway genes are not significantly affected by Pet1 deficiency (Fig 4h-i). These findings indicate that Pet1 controls the chromatin accessibility of CREs of 5-HT neurotransmission genes.

Interestingly, the enhancers associated with *Slc6a4* that are directly occupied by Pet1 show little accessibility to transposase at E11.5 but gradually gain accessibility from E11.5 to E17.5 in wildtype *Pet1* neurons (Fig 4i). In contrast, these TAC regions fail to fully open in *Pet1-/-* neurons (Fig 4i), strongly suggesting Pet1 is directly required for initial conversion of these regions to a euchromatic state. Among the Pet1-regulated TACs, we identified 47 regions that similarly fail to fully open between E11.5 and E17.5 in *Pet1*-/- neurons, 20 of which show direct Pet1 occupancy at E14.5 (Supplemental Fig 5g-h). TACs that require Pet1 for chromatin opening are linked to genes enriched for synaptic products, suggesting that Pet1 opens condensed chromatin at CREs controlling synapse functions (Supplemental Fig 5g).

### Continued Pet1 function is required to sustain 5-HT regulatory element accessibility

Pioneer factors in dividing cells induce “epigenetic memory” such that once “opened”, pioneered sites preserve their accessible state through further rounds of cell division even in the absence of the pioneer factor (Mayran and Drouin, 2018; Pataskar et al., 2016). To determine whether Pet1 is required to sustain stable postmitotic chromatin accessibility, we analyzed flow sorted E14.5 Pet1 conditional knockout (*Pet1^fl/fl; Pet1-Cre; Ai9^*) *Pet1*-lineage neurons (“Pet1-cKO neurons”) in which *Pet1* transcript level is maintained for about 2 days before it becomes lost at E12.5 (Liu et al., 2010) (Supplemental Fig 5i-k). We found that in Pet1-cKO neurons, many 5-HT neurotransmission gene TACs lose accessibility as in *Pet1-/-* neurons (Fig 5a-c) at distal CREs (Fig 5d), including the enhancers of 5-HT neurotransmission genes *Tph2*, *Gch1*, *Ddc*, and *Slc22a3* (Fig 5e-f).

**Figure 5:**
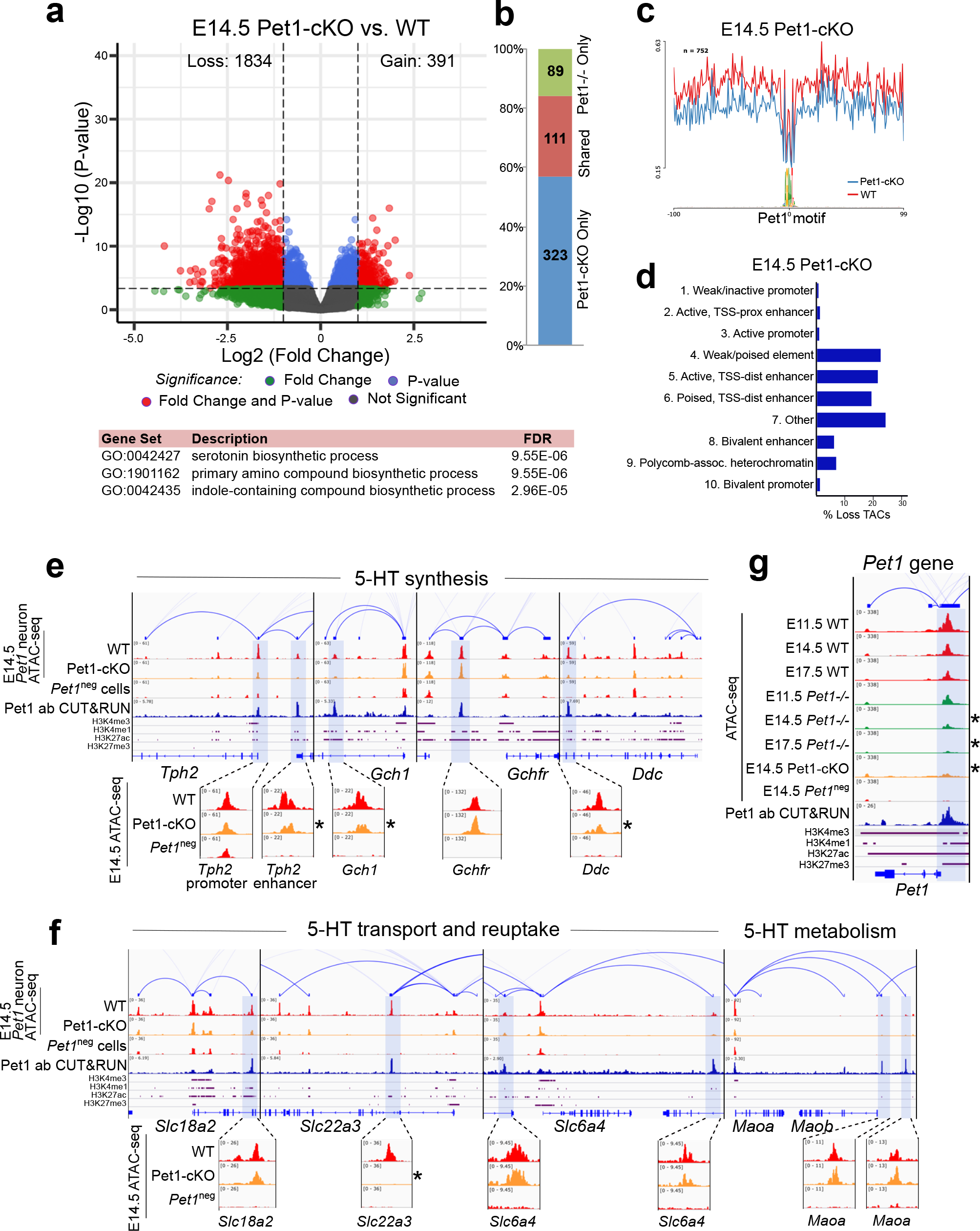
Sustained Pet1 input is required for chromatin accessibility. (a) Volcano plot showing the differential chromatin accessibility of TACs between Pet1- cKO and wildtype *Pet1* neurons, with fold change cutoff of 2-fold and significance cutoff of FDR <0.01 (top). Highest FDR GO terms for the Pet1-maintained TACs (in red) that are occupied by Pet1 show enrichment for serotonin biosynthesis (bottom). (b) Overlap between differential TACs in *Pet1-/-* and Pet1-cKO 5-HT neurons. (c) Comparison of *in vivo* Pet1 footprints over representative ETS-factor motif between Pet1-cKO and wildtype 5-HT precursors within all *Pet1*/5-HT-lineage accessible chromatin regions. (d) The ChromHMM chromatin states genomic coverage for TACs that are maintained by Pet1 at E14.5. (e-f) Genome browser tracks showing the gene-enhancer pairings and the ATAC-seq, Pet1 CUT&RUN, and histone modification ChIPmentation signals at 5-HT identity genes. Zoomed-in inserts display the changes in chromatin accessibility between wildtype (red) and Pet1-cKO (yellow) 5-HT precursors within the highlighted (blue) regions at E14.5. Asterisks denote FDR<0.01 and fold change >2 compared to wildtype. (g) Genome browser tracks showing the ATAC-seq, Pet1 CUT&RUN, and histone modification ChIPmentation signals at the *Pet1* gene locus.

In addition to selecting 5-HT identity genes for transcriptional activation, Pet1 maintains its expression through positive autoregulation (Scott et al., 2005a). CUT&RUN showed that Pet1 directly occupies the TAC that overlaps with its auto-regulatory enhancer (Scott et al., 2005a, 2005b), which is consistent with our previous Pet1 ChIP-seq findings (Wyler et al., 2016) (Fig 5g; Supplemental Fig 5e). We found that Pet1 is required for optimal accessibility of this enhancer region and for sustaining its accessibility suggesting that control of chromatin accessibility is an important mechanism for terminal selector autoregulation (Fig 5g). Together, these findings suggest the early euchromatin landscape in postmitotic 5-HT neurons is a reversible event that requires continuous Pet1 activity for maintenance of accessibility.

### Lmx1b co-regulates reorganization of enhancer chromatin accessibility of 5-HT identity genes

To determine whether the capacity to regulate chromatin accessibility in postmitotic neurons is unique to Pet1, we assessed whether Lmx1b also controls chromatin architecture. Since Lmx1b occupancy in the brain has not been defined, we performed Lmx1b CUT&RUN in E14.5 *Pet1* neurons, which revealed that a significant portion of Pet1 and Lmx1b binding sites coincide within the same TAC regions (Supplemental Fig 6a-c). In Lmx1b-cKO (*Lmx1b^fl/fl; Pet1-Cre; Ai9^*) *Pet1* neurons (Donovan et al., 2019) (Supplemental Fig 6d-f), chromatin landscape is altered (Fig 6a), Lmx1b footprints are reduced (Fig 6b), and more TACs are differentially accessible than in Pet1- cKO neurons (Figs 5a and 6c), suggesting that Lmx1b has a broader impact than Pet1 on 5-HT chromatin. Lmx1b directly binds 19% of the differentially accessible TACs in Lmx1b-cKO (comparable to Pet1 which binds 20% of differentially accessible TACs in Pet1-cKO) indicating direct maintenance of accessibility by Lmx1b at those regions (Fig 6d; Supplemental Fig 5l), which are predominantly distal enhancers (Fig 6e). The Lmx1b-dependent TACs with direct Lmx1b occupancy, of which 65% are specific to *Pet1* neurons (Fig 6h), are also most significantly associated with monoamine/serotonin synthesis (Fig 6c). Indeed, the conditional ablation of Lmx1b results in the even greater closing of 5-HT neurotransmission gene TACs than *Pet1* ablation (Fig 6i-j). Many TACs whose accessibility was not significantly reduced in Pet1-cKO neurons (fold change>2, FDR<0.01) significantly lose accessibility in the Lmx1b-cKO neurons (Supplemental Fig 6i), including the TACs of *Gchfr*, *Slc18a2*, and *Maoa* (Fig 6i-j). Lmx1b directly occupies the enhancers of many 5-HT neurotransmission genes (Fig 6i-j) whose expression is known to be critically dependent on Lmx1b (Ding et al., 2003; Donovan et al., 2019).

**Figure 6:**
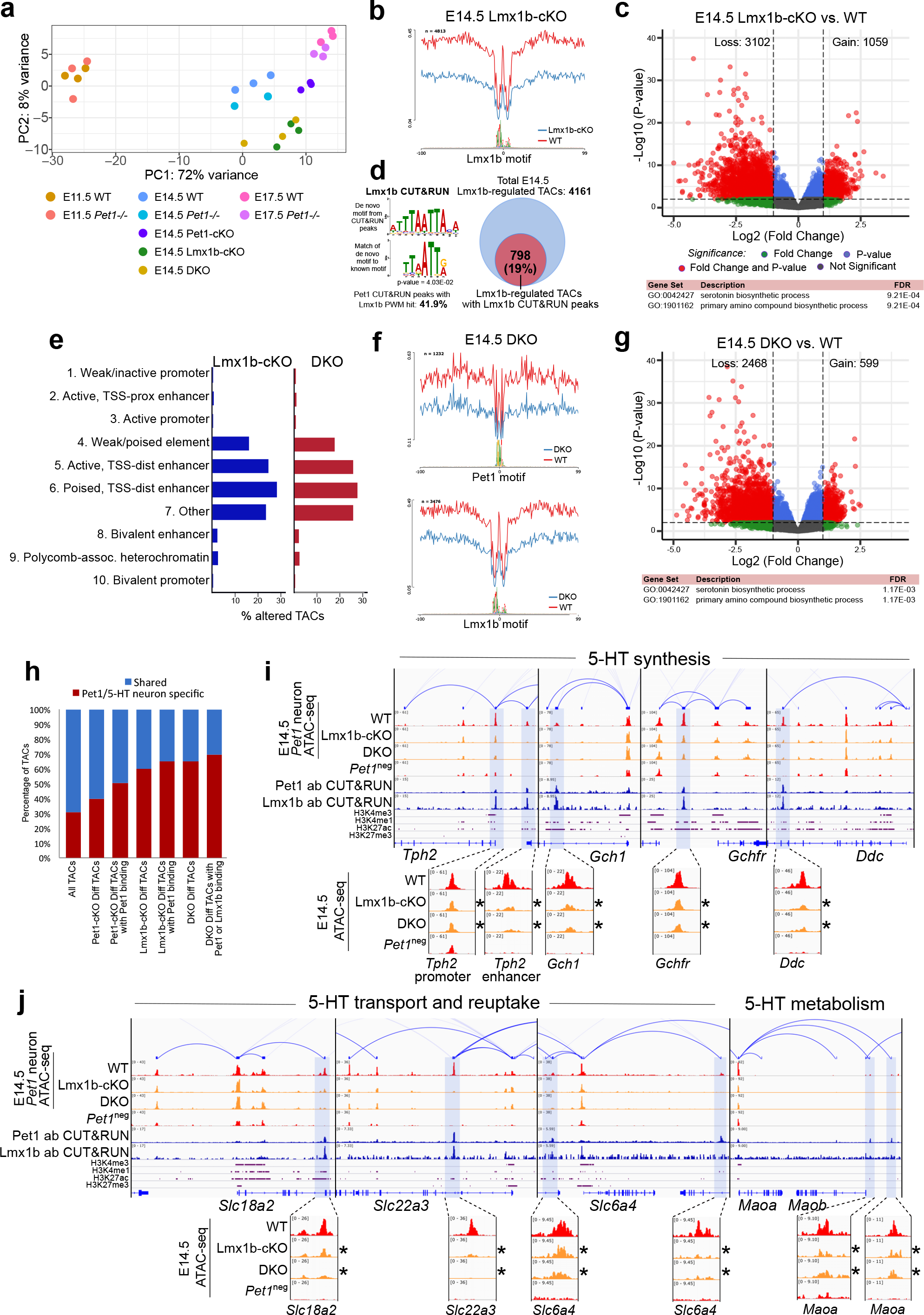
Lmx1b co-regulates chromatin accessibility with Pet1 at 5-HT neurotransmission genes but has broader impact on accessibility landscape. (a) PCA showing the distribution of wildtype, *Pet1-/-*, Pet1-cKO, Lmx1b-cKO, and DKO neuron ATAC-seq data over two principal components. (b) Comparison of *in vivo* Lmx1b footprints over representative LIM-homedomain motif, between Lmx1b-cKO and wildtype *Pet1* neurons within accessible chromatin regions. (c) Volcano plot showing the differential chromatin accessibility of TACs between Lmx1b-cKO and wildtype *Pet1* neurons, with fold change cutoff of 2-fold and significance cutoff of FDR <0.01 (top). Highest FDR GO terms for Lmx1b-maintained TACs (in red) that are occupied by Lmx1b show enrichment of serotonin biosynthesis (bottom). (d) The top most significantly enriched motif within Pet1 CUT&RUN peaks as determined by *de novo* MEME motif analysis matches (*p=4.03E-02*) the Lmx1b LIM-homeodomain binding site previously defined *in vitro* (Jolma et al., 2013) based on analysis by the TOMTOM Motif Comparison Tool (left panel). Fraction of Lmx1b-maintained TACs containing Lmx1b CUT&RUN peaks (right panel). (e) The ChromHMM chromatin states genomic coverage for TACs that lose accessibility in Lmx1b-cKO (blue) and DKO (red) in E14.5 *Pet1* neurons. (f) Comparison of the *in vivo* Pet1 (top) and Lmx1b (bottom) footprints between DKO and wildtype *Pet1*/5-HT lineage neurons. (g) Volcano plot showing the differential chromatin accessibility of TACs between DKO and wildtype *Pet1* neurons, with fold change cutoff of 2-fold and significance cutoff of FDR <0.01 (top). Highest FDR GO terms for Pet1- and Lmx1b-maintained TACs (in red) that are occupied by the two TFs show enrichment for serotonin biosynthesis (bottom). (h) The fraction of Pet1-cKO, Lmx1b-cKO, and DKO differentially accessible TACs that is specific to *Pet1* lineage. A significant fraction of the Pet1- and Lmx1b-regulated TACs is unique to *Pet1* neurons and absent from *Pet1*^neg^ cells. (i-j) Genome browser tracks showing the gene-enhancer pairings and the ATAC-seq, Pet1 CUT&RUN, and histone modification ChIPmentation signals at 5-HT identity genes. Display data ranges for the zoomed-out panels are scaled to the peak in the blue highlighted regions. Zoomed-in inserts display the changes in chromatin accessibility between wildtype (red), Lmx1b-cKO (yellow), and DKO (yellow) 5-HT precursors within the highlighted (blue) regions at E14.5. Asterisks denote FDR<0.01 and fold change >2 compared to wildtype.

We also found that the loss of chromatin accessibility in Lmx1b-cKO is highly similar to that of DKO (*Lmx1b^fl/fl^; Pet1^fl/fl; Pet1-Cre; Ai9^*) *Pet1* neurons (Figs 6b-c and 6f-g), suggesting that the ablation of both TFs does not have greater effect than ablation of Lmx1b alone. PCA also indicates close clustering of Lmx1b-cKO neurons with DKO neurons (Supplemental Fig 6g). These findings are consistent with the model that Lmx1b is required for Pet1-dependent chromatin regulation. At a substantial subset of Lmx1b- dependent TACs such as those of *Tph2*, *Gch1*, *Ddc*, and *Slc6a4* (Fig 5e-f and 6i-j), Pet1 and Lmx1b non-redundantly co-regulate the accessibility of the CREs. In parallel, Lmx1b further sustains, independent of Pet1, an additional subset of TACs that are required for axon and synaptic formation, neurotrophin signaling, and neuronal growth (Supplemental Fig 6h-i).

Motif analysis on the differentially accessible regions in the cKO and DKO neurons showed that the binding sites of several TF families such as the Rfx-related factors are less accessible in Pet1- and Lmx1b-deficient *Pet1* neurons, demonstrating that together Pet1 and Lmx1b impact the global landscape of available TF binding sites (Supplemental Fig 6j).

### Pet1 and Lmx1b regulate 5-HT synaptogenesis

In addition to controlling TACs associated with 5-HT transmission genes that define 5-HT transmitter identity (Fig 6g), Pet1 and Lmx1b program the CRE accessibility of many genes involved in axon growth and synapse formation (Fig 7a; Supplemental Fig 7a), the majority (59%) of which are the same TACs that gain accessibility in wildtype *Pet1* neurons during maturation (Fig 7b). These include the TACs of *Syt1*, which encodes a calcium sensor protein that mediates the release of synaptic vesicles in response to calcium (Yoshihara and Littleton, 2002), and *Slc17a8*, which encodes a vesicular glutamate transporter (vGlut3) that enables glutamatergic co-transmission in a subset of adult 5-HT neurons (Vigneault et al., 2015) (Fig 7c). The Pet1- and Lmx1b- dependent TACs of *Syt1* and *Slc17a8* are *Pet1* neuron specific, suggesting that the expression of pan-neuronal genes are regulated in different neurons by neuron-type specific CREs (Fig 7c). In support of the contribution of these Pet1- and Lmx1b- regulated enhancers to developmental gene expression, *Syt1* and *Slc17a8* expressions are decreased in Pet1-cKO, Lmx1b-cKO, and DKO neurons (Fig 7d) together with the expression of other synaptic and axonal genes linked to Pet1- and Lmx1b-regulated TAC regions (Fig 7e; Supplemental Fig 7b-f). Together these data suggest that in addition to opening enhancers necessary for the acquisition of 5-HT-type transmitter identity, Pet1 and Lmx1b modulate the maturation of accessible CREs for other 5-HT functional properties.

**Figure 7:**
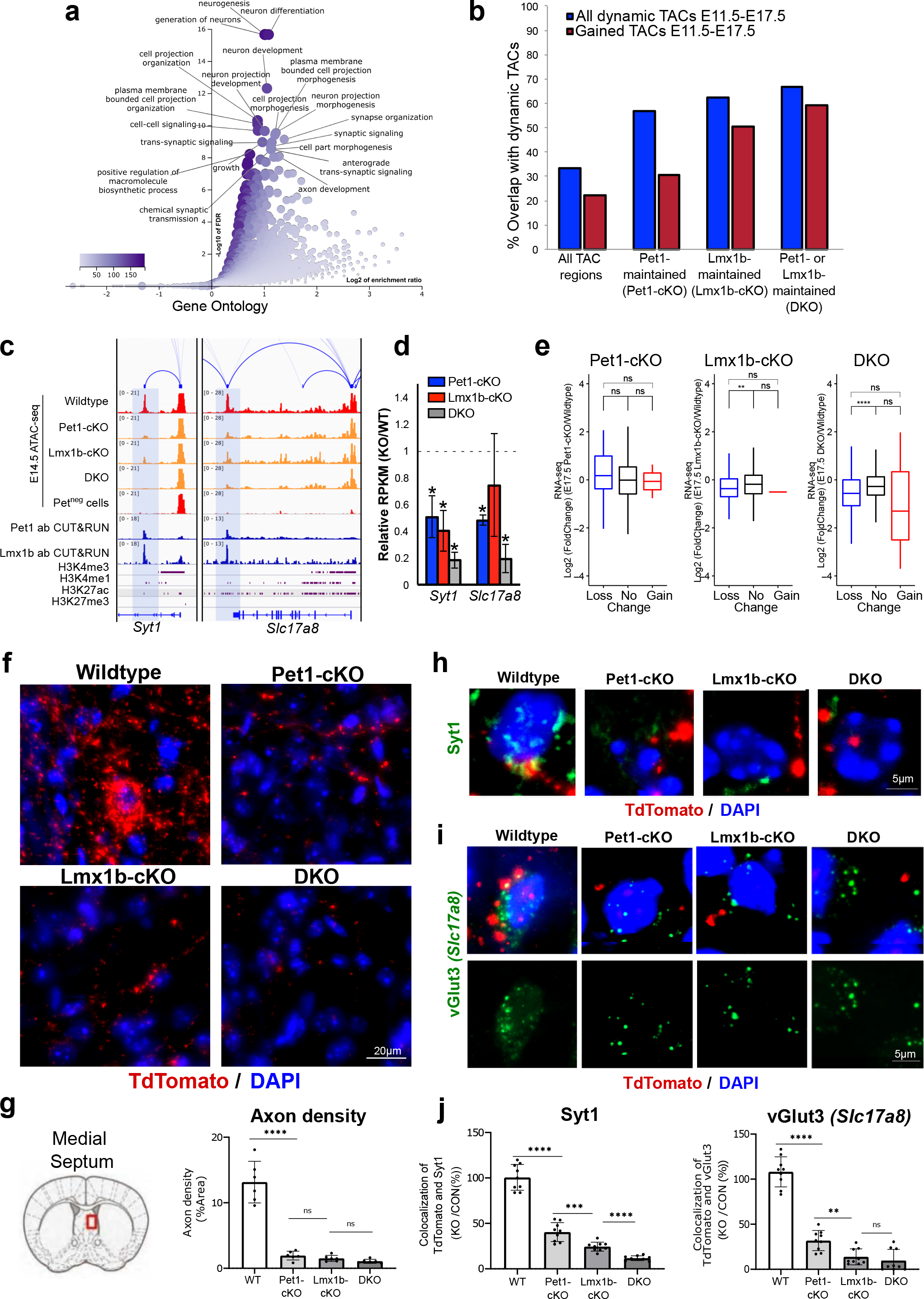
Loss of Pet1 and Lmx1b alters chromatin accessibility at synaptic gene enhancers and disrupts 5-HT neuron connectivity. (a) GO analysis of all Pet1- and Lmx1b-maintained TAC regions, depicted as volcano plot of enrichment (x-axis) vs. FDR statistical significance (y-axis), showing many neuronal biological processes among the top enriched annotations. (b) Fraction of the Pet1- and/or Lmx1b-maintained TACs that change (blue) or gain (red) in accessibility in wildtype *Pet1* neurons from E11.5 to E17.5. (c) Genome browser tracks showing the gene-enhancer pairings and the ATAC-seq, Pet1 and Lmx1b CUT&RUN, and histone modification ChIPmentation signals at synaptic genes *Syt1* and *Slc17a8* show changes in chromatin accessibility between wildtype (red), Pet1-cKO (yellow), Lmx1b-cKO (yellow), and DKO (yellow) *Pet1* neurons within the highlighted (blue) regions at E14.5. Display data ranges for the zoomed-out panels are optimized based on the blue highlighted regions. (d) Relative expression (FPKMs) of *Syt1* and *Slc17a8* in rostral Pet1-cKO, Lmx1b-cKO, and DKO neurons at E17.5. Data presented as mean ± SEM. (e) Differential expression (from E11.5 to E17.5) of the genes associated with Pet1- occupied TACs in Pet1-cKO vs. wildtype 5-HT neurons (left), genes associated with Pet1-occupied TACs in Lmx1b-cKO vs. wildtype 5-HT neurons (middle), and genes associated with Pet1 and Lmx1b co-occupied TACs in DKO vs. wildtype 5-HT neurons (right). (f) TdTomato (red) and DAPI (blue) co-staining of pericellular baskets in the medial septum of P15 *Pet1-Cre; Ai9* wildtype, Pet1-cKO, Lmx1b-cKO, and DKO mice. (g) Diagram showing the analyzed brain area and quantification of TdTomato axons (pixels/μm^2^) (Two-way ANOVA; n = three animals per genotype and two quantified brain sections per animal) (bottom panels). Images were generated through automatic stitching of individual 20x images. (h) Representative immunohistochemistry images of synaptotagmin1 (*Syt1*), TdTomato, and DAPI in the medial septum of P15 *Pet1-Cre; Ai9* mice showing the decreased overlap of Syt1 (green) and TdTomato (red) signals in KO animals compared to wildtype, Images were generated through automatic stitching of individual 63x images. (i) Representative immunohistochemistry images of vGlut3 (*Slc17a8*), TdTomato, and DAPI in the medial septum of adult *Pet1-Cre; Ai9* mice showing the decreased overlap of vGlut3 (green) and TdTomato (red) signals in KO animals compared to wildtype. Images were generated through automatic stitching of individual 63x images. (j) Quantification of Syt1 (left) and vGlut3 (right) co-localization with TdTomato in the medial septum as a percentage of the total TdTomato signal. (Two-way ANOVA; n = two animals per genotype and 3-4 quantified images per animal).

To explore the biological significance of these Pet1 and Lmx1b-dependent TAC alterations, we analyzed the integrity of 5-HT pericellular baskets that encase postsynaptic neuronal dendrites and cell bodies in the medial septum (Senft and Dymecki, 2021). By postnatal day 15 (P15), 5-HT pericellular baskets are readily identified in wildtype brains (Fig 7f; Supplemental Fig 7g). In contrast, in Pet1-cKO, Lmx1b-cKO, and DKO animals, characteristic 5-HT synaptic baskets failed to form (Fig 7f; Supplemental Fig 7g), which is likely attributable to the failure of most axonal projections to reach the medial septum (Fig 7f-g). Of the few axonal terminals that did wrap postsynaptic cell bodies, co-localization of Syt1 and vGlut3 immunostaining with TdTomato+ axon terminals is significantly reduced compared to wildtype animals (Fig 7h-j; Supplemental Fig 7h). This suggests that by regulating the accessibility of CREs of axon and synapse genes Pet1 and Lmx1b are required for both the innervation of postsynaptic neurons and the presynaptic machinery that enables 5-HT and glutamatergic co-transmission.

## DISCUSSION

Specialized molecular, morphological, and functional properties of neurons are largely acquired as postmitotic neurons mature (Polleux et al., 2007). How postmitotic epigenetic programs control gene expression trajectories to encode mature neuronal identities is poorly understood (Bradke et al., 2020; Gallegos et al., 2018). Here, we have defined the accessible chromatin landscape and the cis-regulatory architecture of developing *Pet1* neurons that generate mature 5-HT neurons (Hendricks et al., 1999). We show that the cis-regulatory landscape of embryonic *Pet1* neurons is highly dynamic and strongly correlated with temporal expression trajectories as postmitotic *Pet1* neurons acquire serotonergic identities. Our scATAC-seq and scRNA-seq analyses reveal that *Pet1* neuron chromatin landscapes are highly heterogeneous and established early to prefigure and likely encode 5-HT neuron subtypes including those expressing the glutamatergic marker, *Slc17a8*. We further show that Pet1 and Lmx1b play crucial roles in controlling the dynamics of chromatin accessibility to enable serotonergic neurotransmission and synapse formation. Thus, our findings identify key developmental regulators of chromatin accessibility in postmitotic neurons.

We found that distal transcriptional regulatory element accessibility is heterogeneous not only between *Pet1* neurons and other cell types but also within the *Pet1*-lineage neurons. Diversity in regulatory sequence accessibility is predictive of the transcriptomic outcome of single *Pet1* neurons. Furthermore, the temporal maturation of *Pet1* neuron CRE accessibility is highly correlated with the expression trajectories of their target genes as serotonergic functional properties progressively emerge. TACs linked to genes involved in synapse organization and axon development gain accessibility while those associated with cell cycle control and early embryonic morphogenesis lose accessibility. Importantly, the chromatin landscape is more dynamic between E11.5 and E14.5 than between E14.5 and E17.5, suggesting that the early postmitotic period comprises a highly active stage of chromatin maturation. Since *Pet1* neurons have exited the cell cycle, the temporal changes in open chromatin cannot be explained as an altered abundance of *Pet1* neuron subtypes. Rather, chromatin landscape within individual *Pet1* neurons is dynamic during postmitotic maturation.

The regulatory factors controlling chromatin dynamics in postmitotic neurons are not well defined. TFs with known pioneer activity such as Ascl1 that are expressed in serotonergic progenitors are significantly downregulated or undetectable in postmitotic *Pet1* neurons (Aydin et al., 2019; Wyler et al., 2016). Yet our data suggests that new postmitotic euchromatic regions dynamically appear while other euchromatic regions are lost as 5-HT neuron identity is being acquired, suggesting the existence of regulatory factors that shape postmitotic neuronal chromatin. Additionally, several recent studies computationally mined the genome-wide accessible chromatin sequences and identified TF binding sites that are enriched in regions of dynamic chromatin in neurons or brain regions (Gray et al., 2017; Preissl et al., 2018; de la Torre-Ubieta et al., 2018; Trevino et al., 2021). Here we show that targeting of Pet1 and Lmx1b in the early stages of postmitotic maturation dramatically alters *Pet1* neuron accessible chromatin landscape. CUT&RUN assays suggest that Pet1 and Lmx1b directly control the accessibility of 5-HT specific CREs linked to genes encoding terminal effectors of 5-HT identity and neurotransmission. In fact, we provide evidence in support of a role for Pet1 in initiating the conversion to euchromatin of multiple nucleosome-protected cis-regulatory sequences in postmitotic neurons, including enhancers linked to the serotonin transporter gene, *Slc6a4*. Both Pet1 and Lmx1b are further required to maintain 5-HT specific open chromatin regions, suggesting that postmitotic accessible chromatin landscapes are unstable and require continuous terminal selector input at least during early postmitotic maturation. Finally, we show that Pet1 is required for accessibility of its own autoregulated enhancer region that directs 5-HT specific expression. Autoregulation is thought to boost and maintain terminal selector expression (Leyva-Díaz and Hobert, 2019). We found that in the absence of *Pet1* expression, *Pet1* enhancer accessibility is dramatically reduced but is not fully closed. This suggests that Pet1 boosts its own expression through augmenting auto-enhancer accessibility that was previously initiated by upstream pioneers but was insufficient for proper expression levels of Pet1. Based on these multiple lines of evidence, we propose that Pet1 and Lmx1b define a previously unrecognized category of TFs termed “postmitotic neuron chromatin reorganizers” (PnCRs) that act specifically during postmitotic neuron maturation to shape the accessible chromatin landscapes and thereby select enhancers required for acquisition of specialized neuronal identities. Other TFs may also perform similar function in other neuron types (Allaway et al., 2021; Frank et al., 2015; Preissl et al., 2018). Whereas pioneer factors in progenitor cells can drive cell lineage commitment and induce, through transient input, stable “memories” of epigenetic states that are transmitted through further cell divisions (D’Urso and Brickner, 2014), PnCRs function in differentiated postmitotic neurons and may be required continuously to sustain the euchromatin state of CREs controlling terminal effector genes. The mechanism by which Pet1 and Lmx1b program neuronal chromatin is not understood, but could involve the recruitment of chromatin remodelers or direct cooperative displacement of histones (Spitz and Furlong, 2012).

While Lmx1b maintains accessibility of a distinct set of CREs independent of Pet1, chromatin regulation of many *Pet1-*lineage TACs requires direct and non-redundant co-regulation by both TFs, reminiscent of the cooperative activities of TFs and pioneer factors in other cell types (Holmberg and Perlmann, 2012; Mazzoni et al., 2013; Zaret and Carroll, 2011). Because a large number of cognate motif sites can exist in the genome for each individual TF, selective chromatin accessibility driven by collaborative PnCRs may serve to ensure that neuron-type-specific effector genes are restrictively activated only in the presence of multiple positive regulatory inputs. Altogether, our findings unearth a paradigm by which postmitotic neuron TFs not only activate sequence-specific transcription but also co-program the postmitotic neuronal epigenome to select active gene regulatory sites.

In addition to neurotransmitter synthesis, neurons must develop axonal and synaptic connectivity in order to transmit signals (Kratsios et al., 2015; Spencer and Deneris, 2017). Pet1 and Lmx1b reorganize the access to CREs for both 5-HT neurotransmission and synaptic connectivity. As a result, the acquisition of distinct functional properties is developmentally coupled. A growing body of evidence supports the idea that alterations in 5-HT neurotransmission cause behavioral changes related to those that characterize neuropsychiatric disorders such as anxiety and autism (Doan et al., 2019; Suri et al., 2015). As many of these disorders are neurodevelopmental in origin, the study of the chromatin regulatory programs that control 5-HT neuron and other identities may shed light on the pathophysiology of brain disorders at critical stages of brain development.

## ACKNOWLEDGMENTS

This research was supported by NIH grants RO1 MH117643 and RO1 MH062723 to E.S.D., F30 MH122173 and T32 GM007250 to X.L.Z., and the Uehara Memorial Foundation Fellowship (201940009) to N.T. We thank Katherine Lobur for assistance with mouse breeding and Laura Marsland for assisting with immunohistochemistry analysis of Tph2 in embryonic mouse hindbrain. We thank Dr. Polyxeni Philippidou for helpful comments on the manuscript. We thank Dr. Carmen Birchmeier for the anti- Lmx1b antibody. This work was supported by the Cytometry and Microscopy Shared Resource at Case Comprehensive Cancer Center, the CWRU Genomics Core, and the CWRU Applied Functional Genomics Core.

## AUTHOR CONTRIBUTIONS

W.C.S and E.S.D conceived the original project. X.L.Z., W.C.S., and E.S.D, designed the experiments. X.L.Z, W.C.S, N.T performed experiments. W.C.S. and X.L.Z performed data processing and bioinformatics analyses. X.L.Z, W.C.S, N.T, M.M.K, and E.S.D analyzed and interpreted the findings. X.L.Z, W.C.S, and E.S.D wrote the manuscript.

## METHODS

### Animals

Mice were maintained in ventilated cages on a 12h light/dark cycle with access to food and water *ad libitum* with 2-5 mice per cage. The transgenic mouse lines used in this study have been described previously (Donovan et al., 2019; Wyler et al., 2016). Conditional knockout mice (Lmx1b-cKO, Pet1-cKO, DKO) were generated by breeding animals bearing Ai9 Rosa Tomato (Rosa^Tom^; Jackson Labs Stock #: 007909) and either the Lmx1b^fl/fl^ (Zhao et al., 2006), Pet1^fl/fl^ (Liu et al., 2010), or both alleles with animals harboring *Pet1-Cre* (original name ePet-Cre) transgene. *Pet1+/+; Pet1::EYFP* and *Pet1-/-; Pet1::EYFP* mice were generated by crossing *Pet1+/-* animals bearing the *Pet1::EYFP* transgene. The *Pet1::EYFP* transgene expresses Yfp under the same *Pet1* enhancer as the *Pet1* enhancer region driving the *Pet1-Cre*. Genotyping was performed using ear tissues with the following primers: Pet-1: 5’- CGGTGGATGTGGAATGTGTGCG-3’, 5’-CGCACTTGGGGGGTCATTATCAC-3’, 5’-GCCTGATGTTCAAGGAAGACCTCGG-3’; floxed Pet-1: 5’- TAGGAGGGTCTGGTGTCTGG-3’, 5’-GCGTCCTTGTGTGTAGCAGA-3’; Pet-EYFP: 5’- TATATCATGGCCGACAAGCAG-3’, 5’-GAACTCCAGCAGGACCATGT-3’; floxed Lmx1b: 5’- AGGCTCCATCCATTCTTCTC-3’, 5’- AGGCTCCATCCATTCTTCTC-3’; Rosa TdTomato: 5’-CCACAATAAGCAAGAGGCAC-3’, 5’- CCGAAAATCTGTGGGAAG TC-3’, 5’- CCGAAAATCTGTGGGAAG TC-3’, 5’- CTGTTCCTGTACGGCATG G-3’; Pet1-Cre:5’- CGGCATCAACGTTTTCTTTT -3’, 5’- AGTCAGGGCAGAGCCATCTA -3’.

Embryonic age was defined as the number of days following the appearance of a vaginal plug, designated as embryonic day (E) 0.5. For all procedures, animals were euthanized by anesthesia with isoflurane inhalation followed by cervical dislocation, and brain tissues were collected equally from male and female mice or embryos. All animal experiments were approved by the Institutional Animal Care and Use Committee of CWRU.

### Neuron dissociation and flow cytometry

Isolation of *Pet1* neurons was performed as previously described with a few modifications (Donovan et al., 2019). Briefly, E11.5 or E14.5 embryos from *Pet1- Cre^+^;Ai9* or *Pet1::EYFP+* animals were dissected in cold PBS under dissection microscope to collect all fluorescent cells in the fetal mouse hindbrain. Dissected tissues were transferred into cold 1X TrypLE Express (Gibco) and incubated at 37°C for 15 mins with gentle agitation. Following enzymatic trypsin digestion, samples were centrifuged for 1 min at 1500 rpm, TrypLE Express was removed, and samples were washed and resuspended in 500μL cold Leibovitz’s L-15 media (Thermo Fisher) with 4μL DNAse I (Invitrogen) and 10% FBS (Gibco). Cells were then slowly triturated with fire-polished glass Pasteur pipettes of decreasing bore size until a single cell suspension was achieved.

E17.5 embryos were dissociated using a modified adult neuron dissociation method. *Pet1-Cre^+^;Ai9* or *Pet1::EYFP+* embryos were dissected in cold aCSF solution (3.5mM KCl, 126mM NaCl, 20mM NaHCO3, 20μM Dextrose, 1.25 mM NaH_2_PO_4_, 2mM CaCl_2_, 2mM MgCl_2_, 50μM AP-V (Tocris), 29μM DNQX (Sigma), and 100nM TTX (Abcam)) to collect all fluorescent cells in the hindbrain. Dissected tissues were then incubated in bubbling (95% O_2_, 5% CO_2_) aCSF containing 1mg/mL protease from Streptomyces griseus (Sigma) for 15 mins at room temperature. Following enzymatic digestion, tissues were transferred to 500μL cold aCSF/10%FBS solution containing 4μL DNase I (Invitrogen) and gently triturated with fire-polished glass pipettes to generate single cell suspension.

Dissociated cells were filtered through a 40μm filter (BD Biosciences) and sorted on a Becton Dickinson FACS Aria or FACS Aria-SORP digital cell sorter with 85μm nozzle. Viability of flow sorted cells were routinely checked with Trypan blue staining followed by manual counting of intact cells on small portions of the FACS purified neurons.

### Bulk *Pet1* neuron ATAC-sequencing library preparation

ATAC-seq was carried out as previously described (Buenrostro et al., 2015) with a few modifications using rostral and caudal *Pet1* neurons flow sorted with either *Pet1::EYFP* (wildtype and *Pet1-/-* samples) or *Pet1-Cre;Ai9* (Pet1-cKO, Lmx1b-cKO, and DKO samples). Bulk ATAC-seq for each time-point/condition was performed with 3-5 biological replicates. Each biological replicate consisted of embryos flow sorted on the same day from either one or multiple dams that were then pooled to collect a total of 10,000-50,000 *Pet1* neurons. Neurons were sorted into L15/10%FBS solution, washed with ice cold PBS, and resuspended in 25μL ice cold ATAC-RSB buffer (10mM Tris-HCl, pH 7.4, 10mM NaCl, 3mM MgCl2, 0.1% NP40 (Roche cat#11332473001)). Cells were incubated in ATAC-RSB on ice for 2 mins, then centrifuged at 800 xg for 10 mins at 4°C. Pelleted nuclei were resuspended in 25μL Transposition mix (12.5μL 2x TD buffer (Illumina), 1.25μL transposase (100nM final) (Illumina), 8.25μL PBS, 3.25μL H_2_O) for E11.5 and E14.5 samples and 15μL of Transposition mix for E17.5 samples. The transposition reactions were incubated at 37°C for 25 mins on a Thermomixer at 1000rpm. The resulting DNA fragments were purified using the DNA Clean and Concentrator-5 Kit (Zymo) and PCR amplified for 13 cycles with Illumina Nextera adapter primers using the NEBNext High Fidelity 2X Master Mix (NEB) with the following PCR program: (1) 5 mins at 72°C (2) 30s at 98°C (3) 10s at 98°C, (4) 30s at 63°C, (5) 1 min at 72°C, and (6) Repeat steps 3-5, 12X. Final PCR products were cleaned using PCRClean Dx beads (Aline Biosciences), assessed for quality on a Bioanalyzer, and sequenced for 75 cycles to an average depth of 29M paired-end reads per sample on an Illumina NextSeq 550.

### ChIP-seq using Tagmentation (ChIPmentation)

*Pet1* neurons (caudal and rostral) were flow sorted from *Pet1::EYFP* embryos at E14.5 into PBS/10% FBS. Neurons were frozen in PBS as cell pellets in -80°C until use. 50,000-300,000 frozen cells from multiple days were then thawed on ice, fixed with 1% formaldehyde in PBS for 10 mins at room temperature followed by quenching with 0.125M glycine, and pooled to generate one biological replicate. Two biological replicates were collected for each antibody. ChIP-seq using Tagmentation (ChIPmentation) was then performed as previously described (Schmidl et al., 2015). Cells were washed once with ice cold PBS and lysed in 130μL Sonication Buffer (10mM Tris-HCl pH 8, 0.25% SDS, 2mM EDTA, 1x protease inhibitor cocktail (Active Motif # 37490), 1mM PMSF). Sonication was conducted in MicroTUBEs (130μL) on a Covaris S2 ultrasonicator until DNA fragments were in the range of 200-700bp. Lysates were diluted 1:1.5 with Equilibrium Buffer (10mM Tris-HCl pH 8, 233mM NaCl, 1.66% Triton X-100, 0.166% sodium deoxycholate, 1mM EDTA, 1x protease inhibitor cocktail (Active Motif)) and incubated with 3μg of antibody on a rotator overnight at 4°C. The following antibodies were used: H3K4me3 (Diagenode C15410003-50), H3K4me1 (Abcam ab8895), H3K27ac (Abcam ab4729), H3K27me3 (Active Motif 39055). The next day, 10μL of Protein A/G Dynabeads blocked overnight with 0.1% BSA in RIPA-LS Buffer (10mM Tris-HCl pH 8, 140mM NaCl, 1mM EDTA, 0.1% SDS, 0.1% sodium deoxycholate, 1% Triton X-100) were transferred to each immunoprecipitated lysate and incubated for 2h at 4°C on a rotator. Following incubation, beads were washed twice with ice cold RIPA-LS Buffer, twice with cold RIPA-HS Buffer (10mM Tris-HCl pH 8, 500mM NaCl, 1mM EDTA, 0.1% SDS, 0.1% sodium deoxycholate, 1% Triton X-100), twice with RIPA-LiCl Buffer (10mM Tris-HCl pH 8, 1mM EDTA, 250mM LiCl, 0.5% NP- 40, 0.5% sodium deoxycholate), and twice with 10mM Tris-HCl pH8. Beads were then resuspended in 25μL of tagmentation reaction mix containing 12.5μL of 2x Tagmentation DNA Buffer (Illumina #15027866) and 1μL of Tagment DNA Enzyme (Illumina #15027865). Tagmentation reaction was incubated at 37°C for 10mins with gentle vortex every 5 mins. Tn5 reaction was inactivated by adding 500μL of ice-cold RIPA-LS Buffer. Beads with washed twice with RIPA-LS Buffer and twice with TE Buffer (10mM Tris-HCl pH 8, 1mM EDTA) and resuspended in 48μL of ChIP Elution Buffer (10mM Tris-HCl pH 8, 5mM EDTA, 300mM NaCl, 0.4% SDS) and 2μL of Proteinase K solution (ThermoFisher). Digestion occurred at 55°C for 1h followed by sample de-crosslinking at 65°C overnight. Supernatant was transferred the next day to a new tube, and Proteinase K digestion was repeated on remaining bead-bound chromatin by resuspending the beads in 1μL Proteinase K in 19μL ChIP elution buffer and eluting for 1h at 55°C. Supernatants from both elutions were combined and purified using DNA Clean and Concentrator-5 Kit (Zymo). Library preparation was performed as described for ATAC-seq. ChIPmentation libraries were analyzed on a Bioanalyzer for quality control and then sequenced with 2×75 paired-end reads per sample on an Illumina NextSeq 550.

### Single cell ATAC-seq library construction

*Pet1* neurons from *Pet1::EYFP* embryos were flow sorted at E14.5 into PBS/10% FBS, washed twice with PBS/0.04%BSA, and resuspended in 100μL freshly prepared Lysis Buffer (10mM Tris-HCl pH 7.4, 10mM NaCl, 3mM MgCl_2_, 0.1% Tween-20, 0.1% IGEPAL). After 3 mins of cell lysis on ice, 1mL of chilled Wash Buffer (10mM Tris-HCl pH 7.4, 10mM NaCl, 3mM MgCl_2_, 1% BSA, 0.1% Tween-20) was added to the reaction and nuclei were pelleted and resuspended in 1X Nuclei Buffer (10x Genomics). Nuclei concentration and quality were determined using Trypan blue staining followed by cell counting and inspection on a hemocytometer to ensure that nuclei were free of nuclear membrane blebbing and that there was complete lysis. Nuclei were then added to a transposition mix to perform the Tn5 transposase reaction. The reaction was incubated at 37°C for 60 mins. Transposed nuclei were then loaded onto the 10x Genomics Chromium controller to perform nuclei encapsulation with barcoded gel beads, followed by library amplification for 13 cycles. After purification, the library was analyzed on an Agilent Fragment Analyzer to determine fragment size. Library was sequenced on Illumina Nextseq 550 Sequencing System to a depth sufficient to yield >50,000 paired-end reads per nuclei.

### Single cell RNA-seq library construction

*Pet1* neurons from *Pet1::EYFP* embryos were flow sorted at E14.5 into PBS/10% FBS. FACS purified neuron viability was confirmed to be >80% by staining a portion of the cells with Trypan blue and counting the number of live cells on a hemocytometer. Single cells were loaded in the 10x Single Cell 3’ Chip at a concentration of 1000 cells/μL. Library was prepared according to 10x Genomics Single Cell Protocols Cell Preparation Guide on the 10x Genomics Chromium controller and sequenced to a depth of over 80,000 reads per cell for ∼2,700 cells on Illumina Nextseq 550 Sequencing System.

### CUT&RUN library preparation

CUT&RUN was performed based on the Epicypher protocol v1.6, which is derived from the method developed by Skene, Henikoff, and Henikoff (2018) (Skene et al., 2018). Briefly, Concanavalin A-coated (ConA) beads were washed twice with CUT&RUN Bead Activation Buffer (20mM HEPES pH 7.9, 10mM KCl, 1mM CaCl_2_, 1mM MnCl_2_). FACS purified *Pet1* neurons were washed twice with Wash Buffer (20mM HEPES pH 7.5, 150mM NaCl, 0.5mM spermidine, protease inhibitors (Roche)) and then incubated with activated ConA beads for 15 mins at room temperature to allow the cells to absorb to the beads. ConA-bead bound cells were incubated overnight in 50μL Antibody Buffer (Wash Buffer, 0.01% digitonin, 2mM EDTA) with 1:100 rabbit anti-FEV (Sigma-Aldrich #HPA067679), guinea pig anti-Lmx1b (Carmen Birchmeier, Max Delbruck Center Berlin), or rabbit monoclonal IgG antibody (Jackson ImmunoResearch Lab 115-007-003). One replicate was collected for each antibody. After overnight incubation, the ConA-bound cells were washed twice with Digitonin Wash Buffer (Wash Buffer, 0.01% digitonin), resuspended in 50μL Digitonin Buffer containing 700ng/mL CUTANA pAG-MNase (Epicypher), and incubated for 10 mins at room temperature. After binding of pAG-MNase, ConA bead bound cells were washed twice with Digitonin Buffer, then resuspended in 50μL Digitonin Buffer containing 2mM CaCl_2_ and incubated at 4°C for 2h on a rotator to induce digestion of the chromatin. Reaction was stopped by the addition of 33μL STOP Buffer to each sample (340mM NaCl, 20mM EDTA, 4mM EGTA, 50μg/mL RNase A (ThermoFisher), 50μg/mL GlycoBlue (ThermoFisher)) and incubated at 37°C for 15 mins to release cleaved chromatin into the supernatant and degrade RNA. DNA was purified from the supernatant using the DNA Clean and Concentrator-5 kit (Zymo). Sequencing library preparation was performed using the NEBNext Ultra II Library Prep Kit for Illumina (NEB) according to the manufacturer’s instructions, with the following modification: following adapter ligation DNA was purified using 1.75x PCRClean Dx beads (Aline Biosciences) and libraries were amplified using the following CUT&RUN specific PCR cycling conditions to enrich for 100-700bp fragments: (1) 45s at 98°C, (2) 15s at 98°C, (3) 10s at 60°C, (4) repeat steps 2-3 for a total of 14X, and (5) 1 min at 72°C. PCRClean Dx beads were used to purify the final libraries, which were then sequenced at 75bp x2 on an Illumina NextSeq 550.

### RNA-seq library preparation

Rostral *Pet1* neurons were dissected from E17.5 DKO (*Pet1-Cre; Pet1^fl/fl^; Lmx1b^fl/fl^; Ai9)* or control *Pet1-Cre; Pet1^+/+^; Lmx1b^+/+^; Ai9* embryos and flow sorted into Trizol LS (Invitrogen) on a Becton Dickonson FACS Aria-SORP sorter. Experiment was performed with three control and two DKO biological replicates. Each biological replicate represented 1000-3000 pool flow sorted neurons from multiple embryos of one or more litters. Total RNA was isolated using chloroform extraction and purified using RNA Clean & Concentrator-5 Kit (Zymo). RNA concentration and quality were measured using Quantifluor RNA system (Promega) and Agilent Fragment Analyzer (Advanced Analytics). DNase I treatment and library construction was performed using the Ovation SoLo RNA-Seq Systems V2 according to manufacturer’s instructions. Final libraries were sequenced on a NextSeq 550 (Illumina) with single-end sequencing for 75 cycles.

### Immunohistochemistry, image acquisition, and processing

Mice were deeply anesthetized by isoflurane inhalation (chamber atmosphere containing 4% isoflurane) and perfused for 20 mins with cold 4% paraformaldehyde (PFA) in PBS. Brains were post-fixed in 4% PFA overnight and transferred to 30% sucrose/PBS overnight. Tissues were then frozen in Optimal Cutting Temperature (OCT) solution and sectioned on a cryostat at 25μm thickness. Tissue sections were mounted on SuperFrost Plus slides (Thermo Fisher Scientific) and dried. For antigen retrieval, sections were heated up in Sodium Citrate Buffer for 5 min in the microwave at low power. Sections were blocked with 10% NGS in PBS-T for 1 hr, followed by blocking of endogenous biotin with Avidin/Biotin Blocking kit (Vector) only for Syt1 and vGlut3 stained samples. Sections were incubated with primary antibody (1:1000) at room temperature overnight, then treated the next day with secondary antibodies for 1hr, followed by visualization (Tph2, GFP, and RFP) or further processing with the VECTASTAIN ABC kit (VECTOR) according to manufacturer’s instructions before visualization. Primary antibodies used were: mouse anti-vGlut3 antibody (Abcam ab134310), mouse anti-Synaptotagmin1 (Sigma, SAB1404433), rabbit anti-Tph2 (Millipore Sigma, ABN60), rabbit anti-RFP (Rockland 600-401-379), and chicken anti-GFP (Abcam, ab13970) Secondary antibodies used were goat anti-mouse, Alexa Fluor 488 or 647 (Invitrogen, A10029 or A28181; 1:500), goat anti-chicken, Alexa Fluor 488 (Invitrogen, A11039 #11039, 1:500), goat anti-rabbit, Alexa Fluor 594 (Invitrogen, A11037, 1:500), and biotinylated goat anti-guinea pig IgG antibody (Vector, BA7000; 1:200).

Stained slides were imaged on an LSM800 confocal microscope (Carl Zeiss). In each experiment two animals are analyzed per genotype. Images were acquired per animal using 20X and 63X objective lens and processed in ImageJ. Signal brightness and contrast were edited across whole images equally among genotypes for all pericellular basket synapse and axon images. Signal intensity calculations were normalized for each section. Two-way ANOVA with Welch’s correction analysis was performed.

### ATAC-seq data processing and analysis

ATAC-seq data were processed using the standardized software pipeline from the ENCODE consortium to perform quality-control, read alignment, and peak calling (Landt et al., 2012). FASTQ files from ATAC-seq reads were mapped to UCSC mm10 with Bowtie v2.3.4.3. Samtools v1.9 was used to remove PCR duplicates and chrM reads. Peak calling was performed using MACS2 with the following parameters: --nomodel -- shift 37 --ext 73 --pval 1e-2 -B --SPMR --callsummits. Then by intersecting called peaks in replicates using BEDTools v2.29.0, we defined replicated peaks as the set of peaks present independently in each replicated peak call set and also called in the pooled set.

### Differential accessibility analysis

A comprehensive set of ATAC-seq peaks (TACs) from control datasets (E11.5, E14.5, and E17.5) was generated to be used as the reference for assessing differential accessibility between control time points and for control to TF knockout comparisons. All control peak sets were merged using 1bp overlap to create the union of all control peaks (E11.5, E14.5, and E17.5). Any potential batch effects were corrected using surrogate variable analysis (SVA) (Leek, 2014). The number of significant surrogate variables was determined using an iteratively re-weighted least squares approach. One significant surrogate variable was found and added to the model design in DESeq2 (Love et al., 2014). Differentially accessible peaks were determined using DESeq2 v1.32.0 with a fold change cutoff of 2-fold and FDR adjusted p-value ≤0.01 for all wildtype and conditional knockout comparisons and ≤0.05 for *Pet1-/-* comparisons. BEDTools v2.30.0 was used to find shared intersections between identified peaks (1bp minimum overlap). Peak annotations were performed using ChIPseeker v1.28.3. Principal component analysis was performed using the prcomp function within the ‘plotPCA’ wrapper in DESeq2 on variance stabilizing transformed count data to correct for the underlying mean-variance relationship and plotted with ggplot2 (Love et al., 2014). Volcano plots were generated using the EnhancedVolcano package in R (v1.10.0, https://github.com/kevinblighe/EnhancedVolcano).

### Motif enrichment

TF motifs were identified using monaLisa v0.1.50 (https://fmicompbio.github.io/monaLisa/index.html). Differentially accessible peak regions were binned into “loss”, “no change”, and “gain” categories and motif enrichment for each bin was compared against an equal number of sequences randomly sampled from the genome matched by GC composition. Statistically significant enriched motifs were determined using a one-tailed Fisher’s exact test with Benjamini-Hochberg adjusted p-values ≤ 1e-8.

### Gene ontology enrichment

Gene ontology was performed using Webgestalt requiring >5 and <2000 genes per category and FDR ≤0.05 (Liao et al., 2019).

### ChIPmentation data processing and analysis

ChIPmentation data were processed using the standard ENCODE ChIP-seq software pipeline from the ENCODE Consortium. Sequencing reads were mapped to the UCSC mm10 by Bowtie v2.3.4.3. PCR duplicates were removed using samtools v1.9. The unique mapped reads were used to call peaks comparing immunoprecipitated chromatin with input chromatin with MACS2 v2.2.4 with FDR adjusted p-value cutoff of 0.01. Peak lists were filtered to remove peaks overlapping ENCODE blacklisted regions. BEDTools was used to find shared intersections between identified peaks (1bp minimum overlap). Peak annotations were performed using ChIPseeker v1.28.3.

### Chromatin state analysis

Chromatin state analysis was performed using ChromHMM v1.22 (Ernst and Kellis, 2017). Four histone mark ChIPmentation datasets were binarized at a resolution of 200 bp with a signal threshold of 4. We trained a 10-state model as it characterized the key interactions between the histone marks. The posterior probability state of each genomic bin was calculated using the learned model.

### Super-Enhancer analysis

The rank-ordering of super-enhancers (ROSE) package was used to identify super-enhancers based on the ranking of H3K27ac signal intensities from E14.5 *Pet1* neurons (Lovén et al., 2013; Whyte et al., 2013). Clusters of E14.5 ATAC-seq peaks within 12.5 kb were grouped and ranked for their total input background subtracted H3K27ac signal with super-enhancers called above the inflection point of 6391.9.

### scATAC-seq analysis

Raw scATAC-seq data was processed using the CellRanger ATAC v1.2.0 pipeline (10X Genomics). Reads were aligned to the GRCm38 (mm10) mouse genome. Peaks were called using MACS2 v2.2.4. The peak/cell matrix and all unique single-cell fragments were imported into R using the Signac package (Stuart et al., 2020). Several QC metrics were calculated to assess the quality of the data. Cells with >3000 to <100000 peak region fragments, >40% reads in peaks, <0.025 blacklist ratio, <4 nucleosome signal, >2 TSS enrichment were retained for further analysis. The data was normalized and dimensionality reduced using latent semantic indexing (LSI) (Cusanovich et al., 2015).

First, term frequency-inverse document frequency (TF-IDF) normalizes across cells for sequencing depth and across peaks to give higher values to rare peaks. Next, we ran singular value decomposition (SVD) on the TF-IDF matrix using all variable features. A gene activity matrix was constructed by counting the number of fragments that map within the 2 kb upstream region of each gene. Motif enrichment and TF activity for scATAC-seq clusters were quantified using the “RunChromVAR” wrapper function within the Signac package (v1.14.0) (Schep et al., 2017; Stuart et al., 2020). The filtered cell (n=1692) by peak (n=124111) matrix was used as input with the curated database of mm10 cisBP motifs (n=884). Default settings were used to compute motif accessibility deviation Z-scores.

### scRNA-seq analysis

Raw scRNA-seq data was processed using the CellRanger v3.1.0 pipeline (10X Genomics). Reads were aligned to the GRCm38 (mm10) mouse genome. The unique molecular index (UMI) count matrix was imported into R using Seurat (Butler et al., 2018; Hao et al., 2021; Satija et al., 2015; Stuart et al., 2019). Cells with < 200 or > 7500 expressed genes or > 10% mitochondrial genes were excluded from further analysis. Gene counts were log normalized, then the dataset was scaled and centered and mitochondrial gene levels were regressed out. Dimensionality of the data was determined using an elbow plot and the first 20 principal components were used to reduce dimensionality using principal component analysis.

### Integration of scATAC-seq and scRNA-seq data

scATAC-seq and scRNA-seq datasets were integrated using SCTransform from Seurat using 3000 variable features. To perform clustering, we first computed a principal components analysis (PCA) on the normalized count matrix and used the first 20 PCs to generate a shared nearest neighbor (SNN) graph with the k parameter set to 20. Then we clustered the data with a resolution of 0.8 using the Louvain algorithm. Next, we performed uniform manifold approximation and projection (UMAP) to project the clustering onto low dimensional space and plot the first two dimensions for visualizations (30 nearest neighbors, minimum distance = 0.3) (McInnes et al., 2018). Cluster markers were determined using the Wilcoxon Rank Sum test with a minimum fraction of cells expressing the gene at 0.25 and a log fold-change threshold of 0.25 for each cluster versus all other clusters. The top two marker genes for each cluster are used to identify each cluster.

### Enhancer-gene association

scATAC-seq and scRNA-seq seurat objects were imported into SnapATAC (Fang et al., 2021). A gene expression profile for each cell based on the weighted average of the nearest neighboring cells (k = 15) was imputed based on the scRNA-seq dataset. From the integrated scATAC-seq and scRNA-seq datasets, a “pseudo” multiomics cell is made that contains both gene expression and chromatin accessibility information. Then, logistic regression was performed to quantify the association between gene expression and the binarized accessibility state at putative enhancer elements in a 500 kb window centered at the TSS of each gene. The gene to putative enhancer associations were filtered with p-value ≤ 0.05.

### RNA-seq analysis

RNA-seq reads were aligned to UCSC mm10 using Hisat2 (v2.1.0) with a penalty of 12 for a non-canonical splice site, maximum and minimum penalty for mismatch of 1,0, and maximum and minimum penalty for soft-clipping of 3,0 parameters (Kim et al., 2019). Gene expression quantification and differential expression were calculated using Cufflinks v2.2.2 with a fold change cutoff of 1.5 fold and FDR cutoff of 0.05 (Roberts et al., 2011; Trapnell et al., 2010, 2012).

### CUT&RUN data processing and analysis

CUT&RUN data were processed using the standard ENCODE ChIP-seq software pipeline from the ENCODE Consortium. Sequencing reads were mapped to the UCSC mm10 by Bowtie v2.3.4.3. PCR duplicates were removed using samtools v1.9. The unique mapped reads were used to call peaks comparing TF-bound chromatin with an IgG control with SEACR using both stringent and relaxed threshold settings (Meers et al., 2019). The stringent threshold uses the peak of the total signal curve and the relaxed threshold uses the total signal of the “knee” to the peak of the total signal curve. Both threshold settings have been shown to have high sensitivity and specificity (Meers et al., 2019). BEDTools was used to find shared intersections between identified peaks (1bp minimum overlap). Peak annotations were performed using ChIPseeker v1.28.3. CUT&RUN data for Pet1 and Lmx1b were normalized together for visualization using the ChIPseqSpikeInFree method to determine the scaling factor for adjusting signal coverage across the genome (Jin et al., 2020).

### Footprinting analysis

Transcription factor footprints were identified using RGT-HINT (Gusmao et al., 2016; Li et al., 2019). First, we searched for motifs from HOCOMOCO v11 within the set of differentially accessible ATAC-seq peaks for each TF knockout condition (Kulakovskiy et al., 2018). Then, differential footprints were called on uniquely-mapped paired-end ATAC-seq reads with bias correction for the Tn5 transposase within the set of differentially accessible ATAC-seq peaks comparing control to each TF knockout condition.

### Genomic visualization

ATAC-seq, ChIPmentation, and CUT&RUN data were visualized in Integrative Genomic Viewer (IGV, v2.9.3). Bam files for all ATAC-seq and ChIPmentation datasets were normalized to 1x genome coverage using deeptools (v3.5.0) bamCoverage.

### Statistical analysis

No statistical methods were used to predetermine sample size. Measurements for all experiments were taken from distinct samples. No methods were used to determine whether the data conformed to the assumptions of the statistical method. Experimenters were blinded to genotype during immunofluorescence analysis of 5-HT pericellular baskets. No other blinding of experiments was performed. Clustering of scATAC-seq and scRNA-seq was performed in an unbiased manner. Cell-subtypes were assigned after unbiased clustering. Nuclei that did not meet quality thresholds were removed from further analyses as described in the results and methods.

### Data visualization

Data plots were generated using ggplot2 (v3.3.5) in R (v4.0.4) (Wickham et al., 2016) and Microsoft Excel. 5-HT neurotransmission diagram was created using Biorender.com.

### Code availability

No custom code was used in this study. Open-source algorithms were used as described in the analysis methods. Details on how these algorithms were used to analyze data are available from the corresponding author upon request.

### Data availability

All data generated in this study are deposited in NCBI GEO under accession code GSE185737.

---

**Supplemental Fig 1, related to Figure 1:**
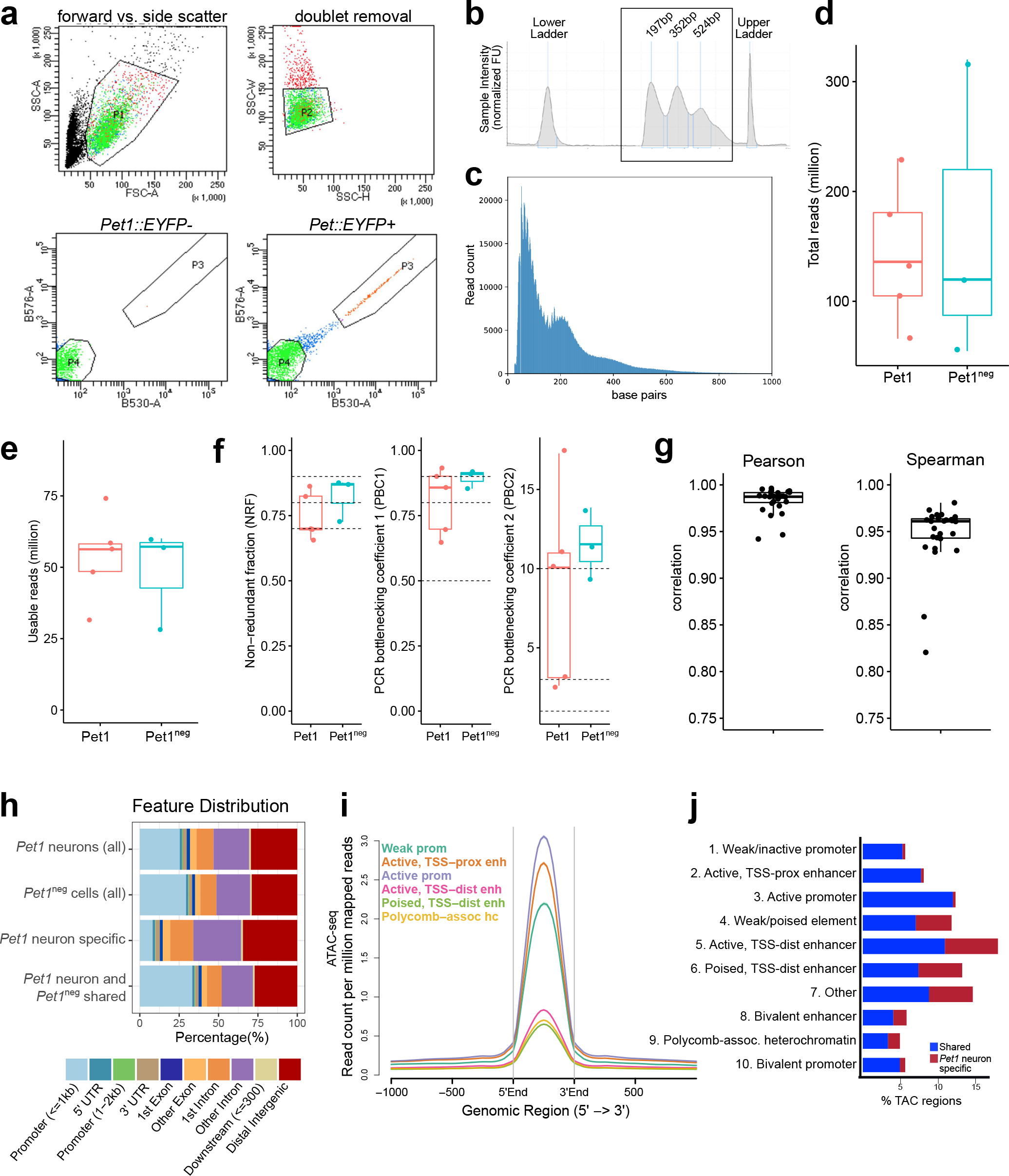
ATAC-seq quality control and analysis. (a) FACS plots of collecting Yfp+ neurons (*Pet1* neurons) from dissociated E14.5 mouse hindbrain cells using forward vs. side-scattered light gating followed by removal of cell doublets. A unique population of cells appear in the hindbrain of *Pet1::EYFP+* mice compared to *Pet1::EYFP-* mice. (b) Tapestation bioanalysis showing the nucleosome laddering of a representative E14.5 ATAC-seq sample using 10,000 FACS-purified *Pet1* neurons. Mono, di, and tri nucleosome sizes (sequencing adapters included) are labeled. (c) Fragment size distribution of ATAC-seq library. (d) The number of total read pairs per cell type. (e) ATAC-seq quality metrics showing the number of usable read pairs per cell type after filtering for mapping quality and PCR duplicates. (f) Library complexity plotted using three ENCODE data standard metrics (NRF, PBC1, PBC2) (Landt et al., 2012). Dotted lines indicate threshold cutoff for very high, high, and medium library complexity. (g) Correlation of ATAC-seq signals at replicated peaks between biological replicated based on Pearson’s correlation coefficient (left) or Spearman’s correlation coefficient (right). (h) Annotation of ATAC-seq peak genomic features for E14.5 *Pet1* neurons and *Pet1*^neg^ hindbrain cells. (i) Average chromatin accessibility at different chromatin states in E14.5 *Pet1* neurons, validating that promoters and enhancers have strong open chromatin signals. (j) Emission probabilities for histone modifications in ten ChromHMM states and the genomic coverage of each chromatin state for E14.5 *Pet1* neuron specific TACs and *Pet1* neuron TACs shared with E14.5 *Pet1*^neg^ hindbrain cells.

**Supplemental Figure 2, related to Figure 2:**
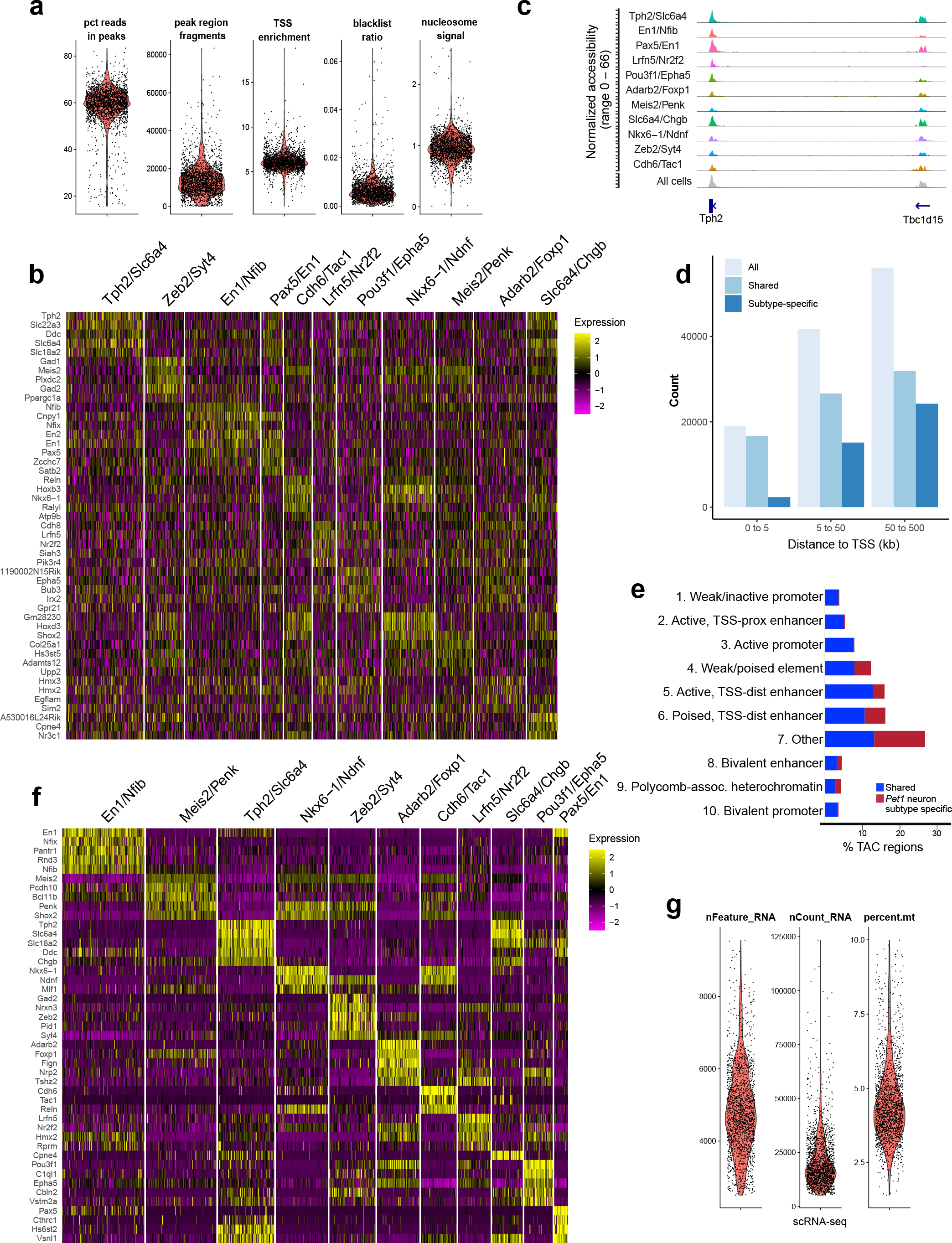
Identification of *Pet1* neuron subtype-specific TACs. (a) scATAC-seq quality metrics showing the distribution of percent of reads in peaks, number of fragments, transcriptional start site enrichment, blacklist ratio, and nucleosome signal per cell. (b) Heatmap showing the marker TACs/chromatin accessibility peaks across scATAC-seq clusters. (c) scATAC-seq tracks showing the aggregated chromatin accessibility peaks at the Tph2 locus for each cluster. (d) scATAC-seq peaks (TACs) sorted by distance to TSS for all TACs, TACs shared by multiple clusters, and TACs that are unique to an individual cluster. A greater fraction of subtype specific TACs are 50-500kb from the TSS than shared TACs. (e) Genomic coverage of ChromHMM chromatin states for scATAC-seq TACs that are specific to a single E14.5 *Pet1* subtype/cluster (red) versus shared among multiple subtypes (blue). (f) Heatmap showing the marker gene expression across scRNA-seq clusters. (g) scRNA-seq quality control metrics showing the distribution of the number of reads, number of genes, and mitochondrial gene fraction per cell.

**Supplemental Figure 3, related to Figure 2:**
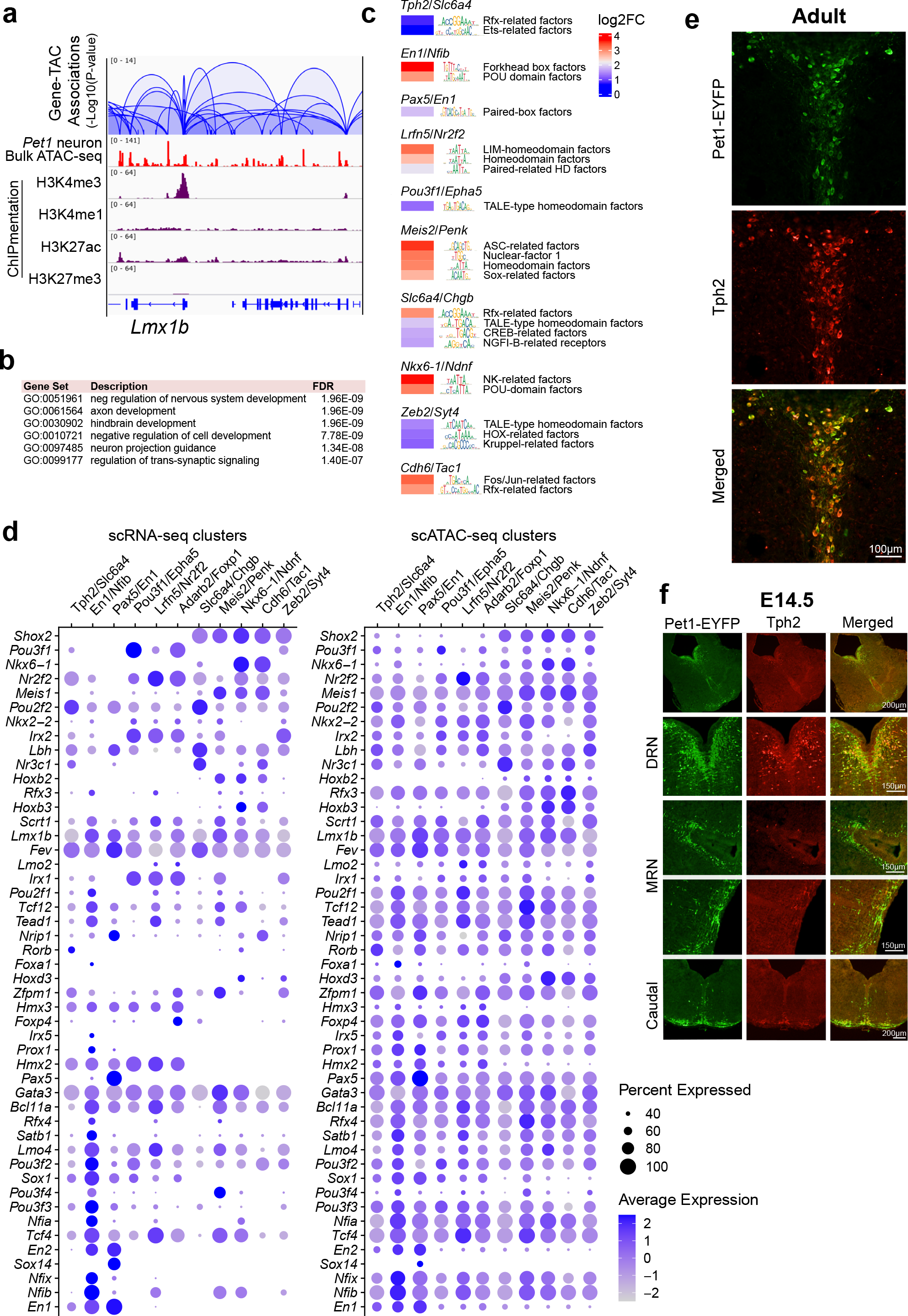
***Pet1* neuron subtype specific TF activity, expression, and accessibility** (a) TACs associated with *Lmx1b*. Loops represent statistically significant TAC-gene association. Loop height represents the p-value of TAC-gene correlation. (b) Top biological functions gene ontology terms for the 1500 genes with the highest number of associated TACs. (c) TF motif enrichment of *Pet1* neuron subtype-specific TACs showing subtype-specific TF activities. (d) Dot plots of gene expression and chromatin accessibility profiles of TFs across *Pet1* neuron subtypes showing a correlation between TF expression and accessibility. Dot size indicates the percentage of cells expressing the gene (left) or exhibiting accessible promoter-proximal chromatin accessibility (right). Color saturation represents the average normalized expression or accessibility level. TFs were selected by their high cluster-dependent variance in expression. (e) Tph2 immunohistochemistry in adult *Pet1:EYFP* mouse brain showing Tph2 and Yfp overlap in dorsal raphe. (f) Tph2 immunohistochemistry in E14.5 *Pet1:EYFP* mouse brain showing partial overlap of Tph2 and Yfp in the dorsal raphe and minimal Tph2 protein levels in the median raphe and within the caudal *Pet1* neurons.

**Supplemental Figure 4, related to Figure 3:**
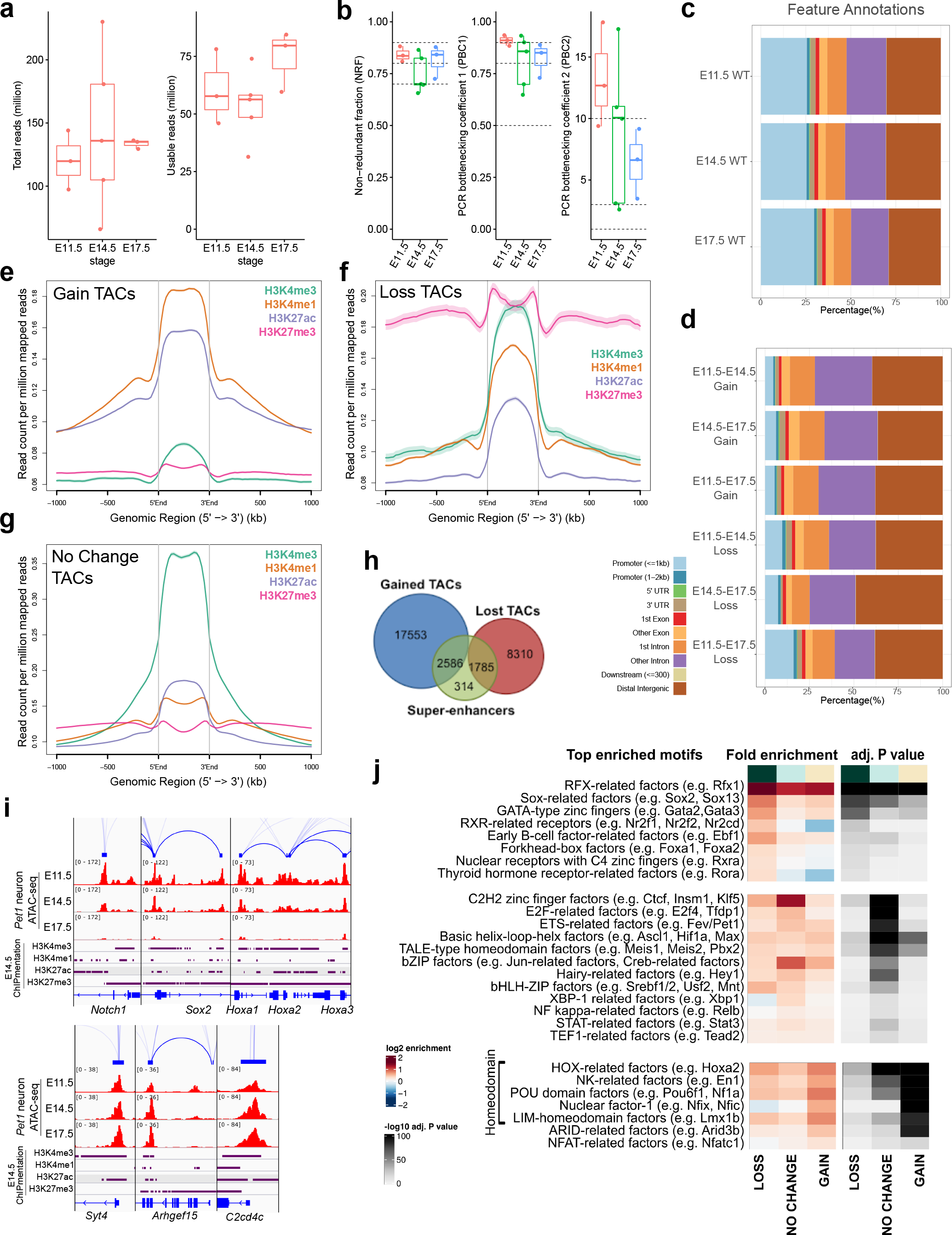
Dynamic developmental remodeling of distal cis-regulatory elements. (a) ATAC-seq quality metrics showing the distribution of total reads (left) and usable reads after filtering for mapping quality and PCR duplicates (right) for each developmental time point. (b) ATAC-seq quality metrics showing library complexity plotted using three ENCODE data standards (NRF, PBC1, PBC2) for each developmental time point (Landt et al., 2012). Dotted lines indicate threshold cutoff for very high, high, and medium library complexity. (c) Distribution of gene feature annotations for all TAC regions at each developmental time point. (d) Distribution of gene feature annotations for TACs that show gain or loss in accessibility between developmental time points. (e) Histone ChIPmentation signals (E14.5) at TACs regions that show gain in chromatin accessibility from E11.5 to E17.5. (f) Histone ChIPmentation signals (E14.5) at TACs regions that show loss in chromatin accessibility from E11.5 to E17.5. (g) Histone ChIPmentation signals (E14.5) at TACs regions that show no change in chromatin accessibility from E11.5 to E17.5. (h) Overlap of all *Pet1* neuron super-enhancers (green) with gained (blue) or lost (red) TACs from E11.5 to E17.5. (i) Genome browser tracks of genes showing loss (left) or gain (right) of chromatin accessibility from E11.5 to E17.5. (j) Motif enrichment within differentially accessible TACs. Highly similar motif hits are combined and assigned to their closest match of a TF class or family. TF families are sorted into three groups by whether their motifs are most significantly enriched in LOSS, NO CHANGE, or GAIN TACs. The TF families are then ranked by fold enrichment within each group. Fold enrichment and p-value are calculated and visualized by the Bioconductor package monaLisa.

**Supplemental Fig 5, related to Figures 4 and 5:**
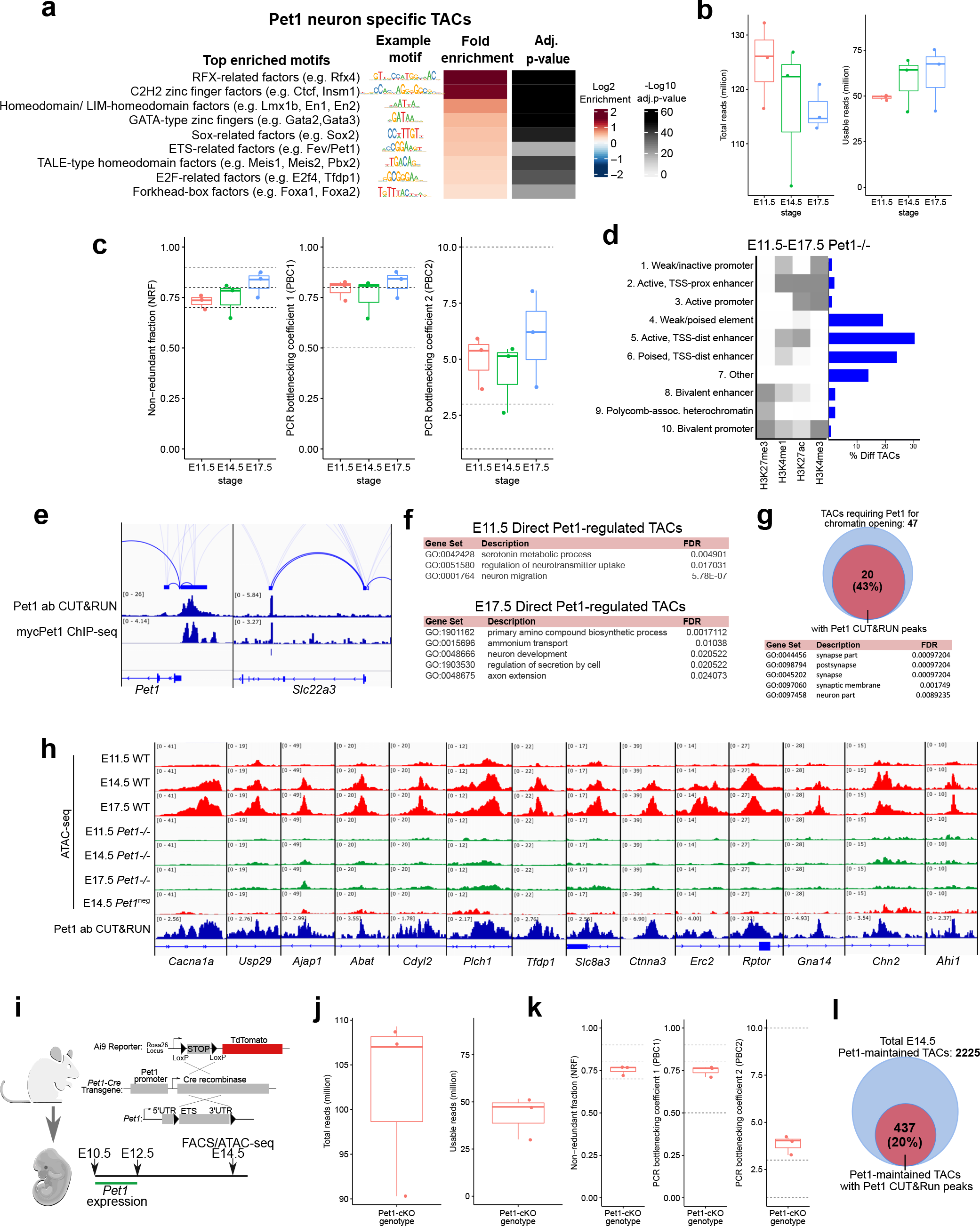
Pet1 is required for chromatin accessibility at *Pet1* neuron specific TACs. (a) Motif enrichment within *Pet1* neuron specific TACs using the Bioconductor package monaLisa. Similar motif hits are combined and assigned to their closest match of a TF class or family. (b) ATAC-seq quality metrics showing the distribution of total reads (left) and usable reads after filtering for mapping quality and PCR duplicates (right) for *Pet1-/-* neurons at E11.5, E14.5, and E17.5. (c) ATAC-seq quality metrics showing library complexity plotted using three ENCODE data standards (NRF, PBC1, PBC2) (Landt et al., 2012) for *Pet1-/-* neurons assayed at E11.5, E14.5, and E17.5. Dotted lines indicate threshold cutoff for very high, high, and medium library complexity. (d) The ChromHMM chromatin states genomic coverage for TACs that lose accessibility in *Pet1-/-* neurons at E11.5, E14.5, or E17.5. (e) Genome browser tracks of Pet1 CUT&RUN and previously published ChIP-seq using myc-Pet1 (Wyler et al., 2016) showing the similarity of Pet1 occupancy peaks at Pet1- regulated genes. (f) GO terms for Pet1-occupied differentially accessible TACs between *Pet1-/-* and wildtype *Pet1* neurons at E11.5 (top) and E17.5 (bottom) show enrichment for serotonin biosynthesis. (g) The fraction of TACs that require Pet1 for chromatin opening from E11.5 to E17.5 that contains Pet1 CUT&RUN peaks (top). Gene ontology analysis on TACs that require Pet1 for initial chromatin accessibility shows enrichment of synapse-related terms (bottom). (h) Representative genome browser tracks of TACs that require Pet1 for euchromatin formation. (i) Schematic of conditional Pet1 targeting and labeling using *Pet1-Cre* transgene followed by FACS collection of TdTomato+ neurons for ATAC-seq. (j) ATAC-seq quality metrics showing the distribution of total reads (left) and usable reads after filtering for mapping quality and PCR duplicates (right) for Pet1-cKO 5-HT precursors at E14.5. (k) ATAC-seq quality metrics showing library complexity plotted using three ENCODE data standards (NRF, PBC1, PBC2) (Landt et al., 2012) for Pet1-cKO 5-HT precursors at E14.5. (l) Fraction of Pet1-maintained TACs (differentially accessible TACs in Pet1-cKO) containing Pet1 CUT&RUN peaks.

**Supplemental Fig 6, related to Figure 6:**
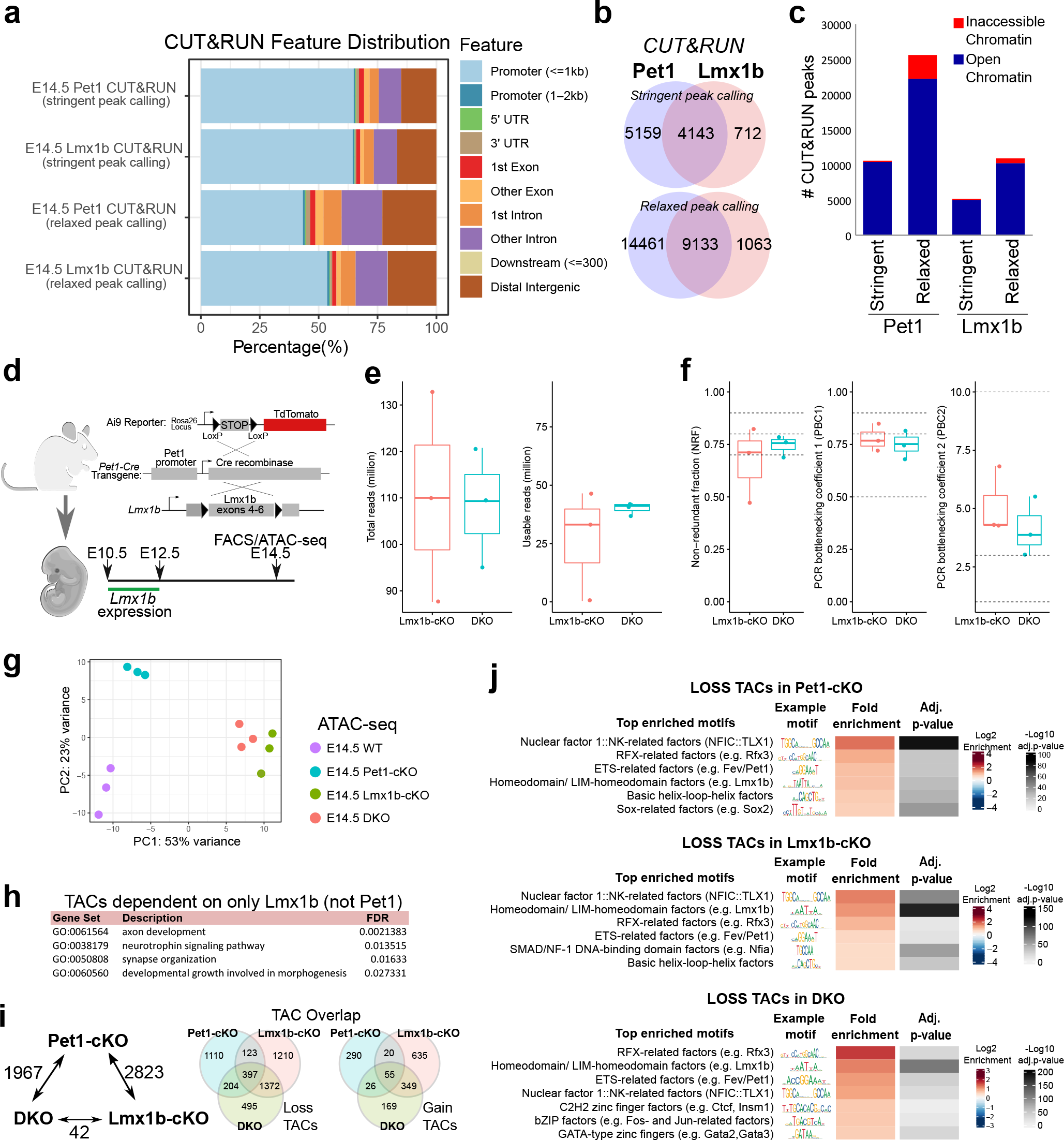
Pet1 and Lmx1b control the global landscape of TF binding sites. (a) Distribution of gene features for E14.5 Pet1 and Lmx1b CUT&RUN peaks using either “stringent” or “relaxed” peak calling parameters (Meers et al., 2019). (b) Overlap of Pet1 and Lmx1b CUT&RUN binding sites defined by either “stringent” or “relaxed” peak selection mode (Meers et al., 2019). (c) Overlap of “stringent” and “relaxed” E14.5 Pet1 and Lmx1b CUT&RUN peaks with E14.5 ATAC-seq TACs showing that Pet1 and Lmx1b mostly occupy open chromatin. (d) Schematic of conditional Lmx1b targeting and labeling using *Pet1-Cre* transgene followed by FACS collection of TdTomato+ neurons for ATAC-seq. (e) ATAC-seq quality metrics showing the distribution of total reads (left) and usable reads after filtering for mapping quality and PCR duplicates (right) for Lmx1b-cKO and DKO at E14.5. (f) ATAC-seq quality metrics showing library complexity plotted using three ENCODE data standards (NRF, PBC1, PBC2) (Landt et al., 2012) for Lmx1b-cKO and DKO 5-HT precursors assayed at E14.5. (g) Principal component analysis showing the distribution of E14.5 wildtype, Pet1-cKO, Lmx1b-cKO, and DKO neuron ATAC-seq data over two principal components. (h) Significant gene ontology terms for the LOSS TACs in Lmx1b-cKO that are not significantly decreased in accessibility in Pet1-cKO. (i) The number of TACs that are differentially accessible between Pet1-cKO, Lmx1b- cKO, and DKO 5-HT neurons by fold change >2, FDR <0.01 (left). Overlap of TACs showing loss (middle) or gain (right) of chromatin accessibility in Pet1-cKO, Lmx1b-cKO, and DKO *Pet1* neurons relative to wildtype. (j) Top enriched motifs within TACs showing loss of accessibility in Pet1-cKO (top), Lmx1b-cKO (middle), or DKO (bottom) neurons relative to wildtype. Highly similar motif hits are combined and assigned to their closest match of a TF class or family. The TF families are then ranked by fold enrichment. Fold enrichment and p-value are calculated and visualized by the Bioconductor package monaLisa.

**Supplemental Fig 7, related to Figure 7:**
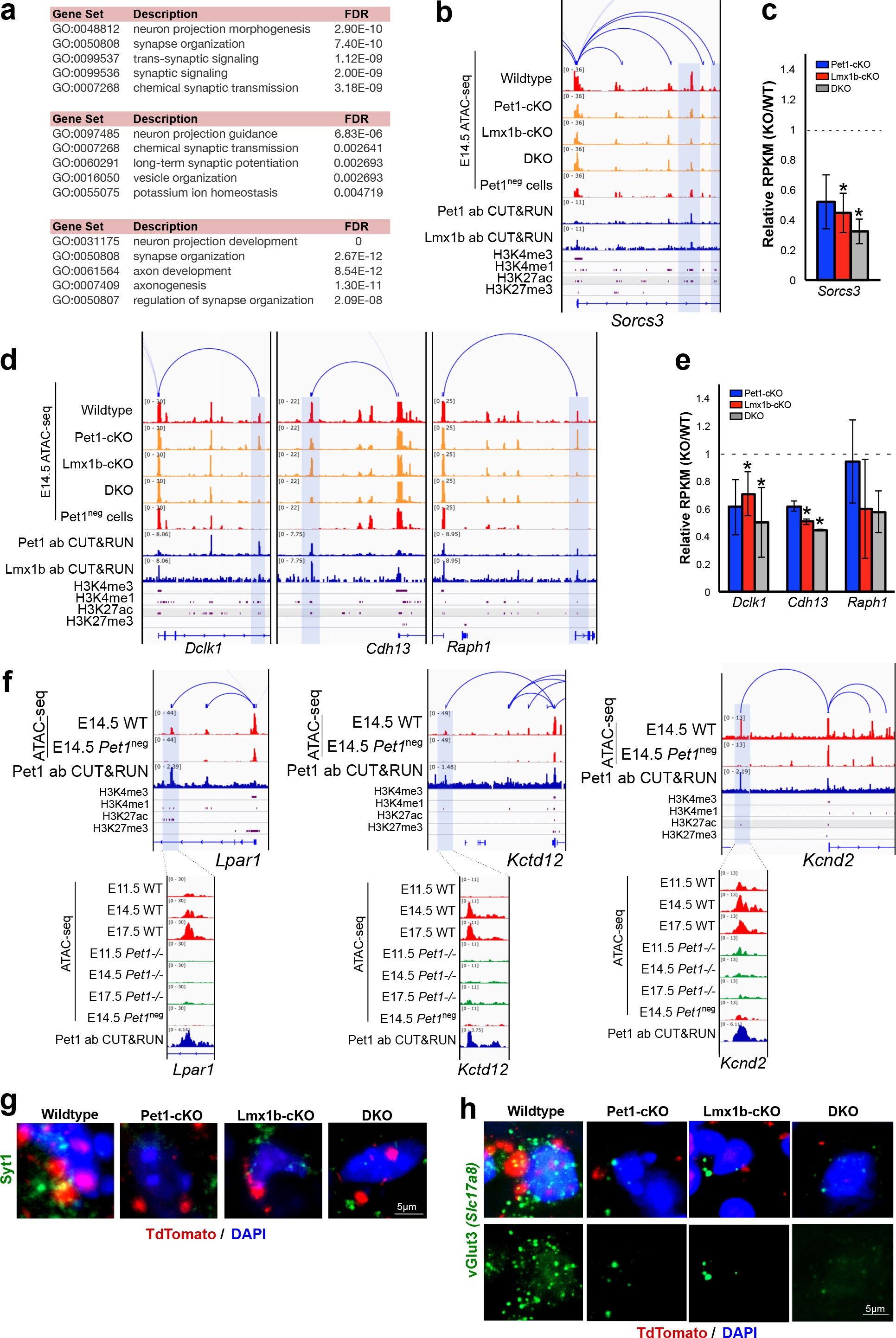
Pet1 and Lmx1b control chromatin access to synapse and axon gene TACs. (a) Top gene ontology terms for the differentially accessible TACs in Pet1-cKO (top), Lmx1b-cKO (middle), and DKO *Pet1* neurons (bottom) ranked by fold enrichment. (b) Genome browser tracks showing the gene-enhancer pairings and the ATAC-seq, Pet1 and Lmx1b CUT&RUN, and histone modification ChIPmentation signals at synaptic gene *Sorcs3.* Display data range is scaled to the peak in the blue highlighted region. (c) Relative expression (FPKMs) of *Sorcs3* in rostral Pet1-cKO, Lmx1b-cKO, and DKO neurons at E17.5. Data presented as mean ± SEM. (d) Genome browser tracks showing the gene-enhancer pairings and the ATAC-seq, Pet1 and Lmx1b CUT&RUN, and histone modification ChIPmentation signals at select axon related genes *Dclk1, Cdh13,* and *Raph1.* Display data ranges are scaled to the peaks in the blue highlighted regions. (e) Relative expression (FPKMs) of *Dclk1, Cdh13,* and *Raph1* in rostral Pet1-cKO, Lmx1b-cKO, and DKO neurons at E17.5. Data presented as mean ± SEM. (f) Genome browser tracks showing the gene-enhancer pairings and the ATAC-seq, Pet1 CUT&RUN, and histone modification ChIPmentation signals at synaptic genes that require Pet1 for chromatin opening. Zoomed-in displays show changes in chromatin accessibility between wildtype (red) and *Pet1-/-* (green) 5-HT precursors within the highlighted (blue) regions across developmental time points. Display data ranges for the top zoomed-out panels are scaled to the peak in the blue highlighted regions. (g) Additional immunohistochemistry images of synaptotagmin1 (*Syt1*), TdTomato, and DAPI in the medial septum of P15 *Pet1-Cre; Ai9* mice showing the decreased overlap of Syt1 (green) and TdTomato (red) signals in KO animals compared to wildtype, Images were generated through automatic stitching of individual 63x images. (h) Additional immunohistochemistry images of vGlut3 (*Slc17a8*), TdTomato, and DAPI in the medial septum of P15 *Pet1-Cre; Ai9* mice showing the decreased overlap of vGlut3 (green) and TdTomato (red) signals in KO animals compared to wildtype, Images were generated through automatic stitching of individual 63x images.

